# Using developmental rules to align microevolution with macroevolution

**DOI:** 10.1101/2022.08.19.504140

**Authors:** Fabio Andrade Machado, Carrie S. Mongle, Graham Slater, Anna Penna, Anna Wisniewski, Anna Soffin, Vitor Dutra, Josef C. Uyeda

## Abstract

Macroevolutionary biologists have classically rejected the notion that higher level patterns of divergence arise through microevolutionary processes acting within populations. For morphology, this consensus partly derives from the inability of quantitative genetics models to correctly predict the behavior of evolutionary processes at the scale of millions of years. Developmental studies (evo-devo) have been proposed to reconcile micro and macroevolution. However, there has been little progress in establishing a formal framework to apply evo-devo models of phenotypic diversification. Here, we reframe this issue by asking if using evo-devo models to quantify biological variation can improve the explanatory power of comparative models, thus helping us bridge the gap between micro- and macroevolution. We test this prediction by evaluating the evolution of primate lower molars in a comprehensive dataset densely sampled across living and extinct taxa. Our results suggest that biologically-informed morphospaces alongside quantitative genetics models allow a seamless transition between the micro and macro scales, while biologically uninformed spaces do not. We show that the adaptive landscape for primate teeth is corridor-like, with changes in morphology within the corridor being nearly neutral. Overall, our framework provides a basis for integrating evo-devo into the modern synthesis, allowing an operational way to evaluate the ultimate causes of macroevolution.

---

“Macroevolution” is the field of study that aims to understand how the diversification of life occurred on our planet over large time scales^1^. Like any other historical science, it seeks to make sense of patterns over time ingrained in the fossil record and phylogenetic trees by referencing well-understood processes known from direct observations and experimentation^2^. In the case of evolutionary biology, this knowledge comes mainly from fields such as ecology and genetics, which tend to map evolutionary phenomena that take place during shorter time scales. For this reason, these studies are sometimes called “microevolution” and are designed to understand how population-level phenomena can produce evolutionary change. However, despite the presumed direct relationship between micro- and macro levels, quantitative studies have struggled to explain most macroevolutionary patterns in terms of microevolutionary processes^3–7^. Nevertheless, empirical results have consistently shown that the availability of additive genetic variation correlates strongly with rates of macroevolution for different traits^6,8–13^, suggesting some effect of lower level microevolutionary processes at the macroevolutionary scale. Whether we can bridge the gap between these two scales is still unclear, with some authors arguing for their essential irreconcilability^3,14,15^ and others advocating for reconciliation within the context of the modern synthesis^5,8–10,16–23^.

One long-standing suggestion for bridging the gap between micro- and macroevolution has been through the study of developmental biology and ontogeny (*i.e.* evo-devo)^10,19,24–28^. This suggestion, however, has been challenging to implement. In a microevolutionary context, development can often be reasonably assumed to be a smooth genotype-to-phenotype (GP) map, *i.e.* genotypic variation translates to phenotypic variation in a linear way, with traits being influenced by multiple genes of smaller effect. Such a smooth GP map would, in turn, allow the modeling of evolution and adaptation of the adult phenotype using a quantitative genetic framework, precisely because these classes of models entail this simplified, linear GP mapping^18–20,29^. On larger time scales, however, genetic architectures can change, selection can fluctuate, and development can be reorganized, generating non-linearities between genotypic and phenotypic divergence, even if the GP map was originally smooth. On the phenotypic level, these non-linearities can produce discontinuities, *i.e.*, regions of the morphospace less inhabited, or not inhabited at all by species, then impeding a straightforward extrapolation of microevolutionary processes over millions of years^19,27,30–32^ Therefore, in the absence of in-depth knowledge of development and the GP map, it is likely that macroevolutionary studies will find heterogeneity and discontinuities, even if the underlying genetic changes at the microevolutionary scale are relatively smooth and continuous^19^. Alternatively, if we can use developmental models as the basis of the quantification of morphology, we might smooth out some of these non-linearities, maximizing our ability to seamlessly connect micro and macroevolutionary scales^9,10,19,30,33–36^.

Here, we propose a new framework for the investigation of morphological evolution over macroevolutionary time that explicitly models evolution at this scale as a consequence of underlying microevolutionary processes (Fig. 1). To deal with potential non-linearities that might arise over long time scales, we suggest the construction of developmentally informed spaces (Fig. 1Bi), which coupled with quantitative genetics modeling (Fig. 1E-G) and comparative methods (Fig. 1D), can facilitate a conceptual bridge between micro and macro scales. Under our proposed framework, we are able to directly compare microevolution-inspired models (henceforth called “microevolutionary models”) to non-microevolution-inspired ones that account for a wider variety of rate and state-heterogenous evolutionary processes (henceforth called “macroevolutionary models”).

**Figure 1.**
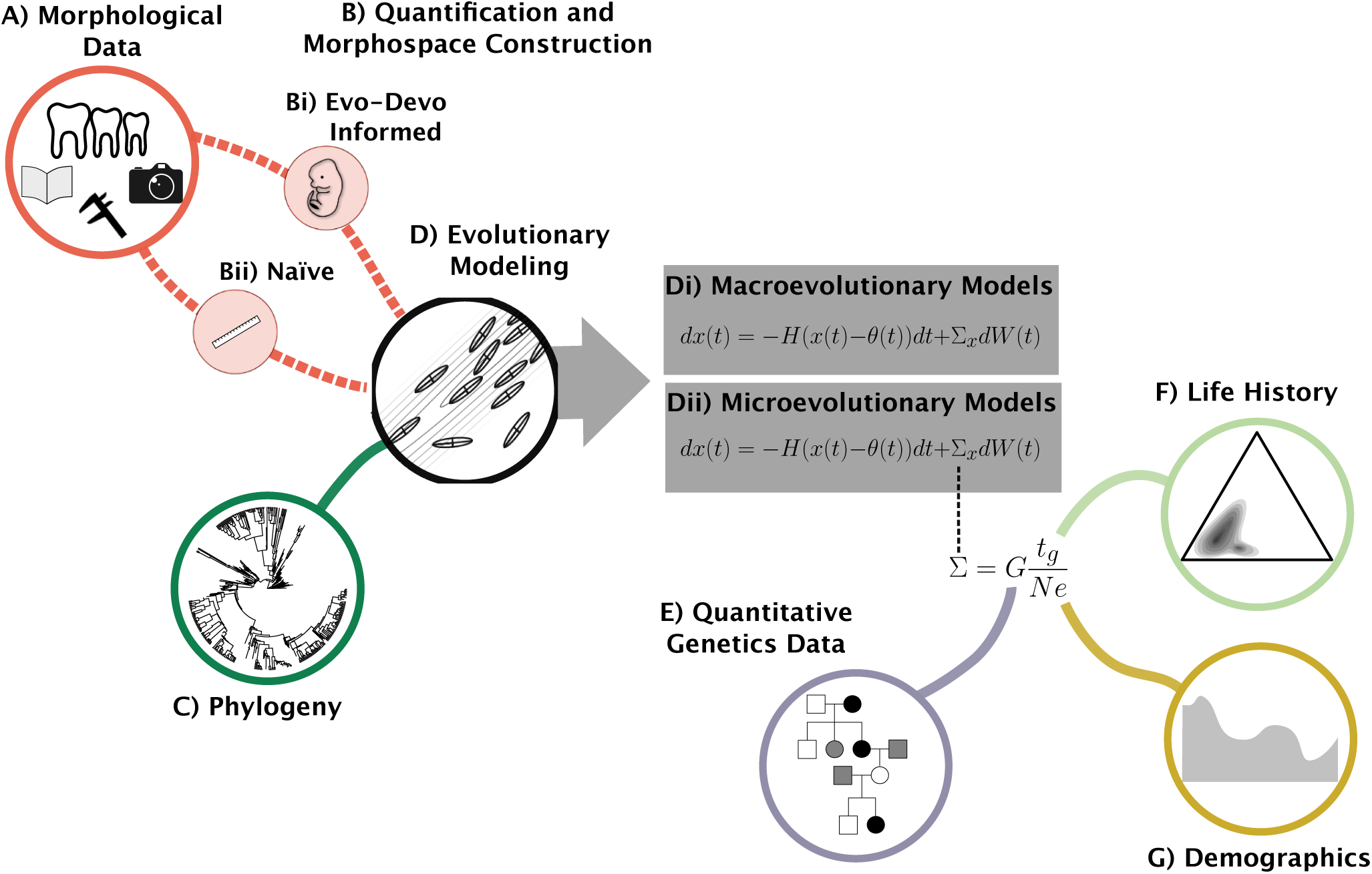
Integrative evolutionary modeling framework employed in the present study. A) Sources of morphological data, such as direct measuruments with calipers, measurements extracted from photographs and data obtained from the literature. B) Process of quantification, which can be either evo-devo inspired (Bi) or “naïve” in relation to developmental processes (Bii). C) Phylogeny of the group under study. D) Evolutionary modeling used to infer adaptive landscapes (isolines) were species (ellipses) have evolved. E) Quantitative genetics data, specifically additive genetic variance-covariance matrices (*G*), estimated form pedegreed populations. F) Life history data, specifically the generation time used to estimate time of divergence in generations (*t_g_*). G) Demographics data, specifically effective population size (*N_e_*). Evolutionary models can be either macroevolutinary (Di) or microevolutionary (Dii). Both models belong to the Orstein-Uhlenbeck class of models with the same parameters (see Materials and Methods for a full description of parameters) with the single difference being that on the microevlutionary models, the stochastic rate parameter Σ is constrained to be equal to the rate of genetic drift. The rate of drift is modeled as being proportional to *Gt_g_/N_e_*). In our framework, the morphological data (A) is used to construct naïve or biologically informed morphospaces (B) and, together with a phylogeny (C), are used in an evolutionary modeling process (D). Different models, including microevolutionary and macroevolutionary ones can then be directly compared when they are estimated under the same morphospace.

We test this workflow to investigate the evolution of primate molars, which is an ideal model system for the present investigation. First, there is a simple yet powerful evo-devo model that describes the development and evolution of Mammalian molars, the inhibitory cascade model (ICM). The ICM models teeth size (*i.e.* molar row form) as the result of a balance between inhibition and activation factors^35^. Specifically, it predicts that the sizes of the first, second and third molars (m1, m2 and m3, respectively) will either be the same (m1=m2=m3), increase (m1<m2<m3) or decrease (m1>m2>m3) along the molar row. A corollary of this prediction is that there will be a positive relationship between the ratios of the areas of the last two molars in relation to the first one (m2/m1 and m3/m1) and thus establishes a natural morphospace to investigate this developmental process. This model was initially proposed for rodents^35^, and later verified for multiple Mammalian species^37,38^, including Primates^11,39,40^. Second, there are several studies characterizing aspects of additive genetic variation in molars for Primates^41–44^, as well as large-scale life history and demographic information for the group^45,46^, parameters that are essential to model microevolutionary processes such as drift and selection (Fig. 1E-G). Third, tooth enamel is the most mineralized substance in vertebrate tissues, making teeth especially resistant to taphonomic processes and abundant in the fossil record (Fig. 1C). The use of a dense fossil record allows us to bridge some phylogenetic gaps between extant species, ensuring that heterogeneities along the tree are more likely due to differences in process rather than incomplete sampling. This extensive availability of paleontological and neontological data enables unprecedented power to evaluate evolutionary dynamics through deep-time using data-hungry phylogenetic comparative methods^47–49^. We apply our framework to an expansive dataset of both extant (232 taxa) and extinct (248 taxa) species summarized from more than 250 different sources (see Supplementary Material), integrated with a newly published comprehensive phylogeny^50^. To address our hypotheses, we use a model-fitting approach based on information theory (Bayesian Information Criteria, or BIC) and model simplicity (minimizing parameter number). We expect that variables devised to quantify developmental processes (ICM variables) will favor microevolutionary models, while data embedded in biologically “naïve” spaces (those with no direct correspondence to any developmental model) will favor complex macroevolutionary models.

## Results

To test our hypothesis that the use of developmental models to quantify morphological variation would provide a better bridge between micro and macroevolution, we used three morphospaces (Fig S1). The first is based on the linear distances taken directly from the teeth (“distance space”), and the second is based on the occlusal areas of each molar (“area space”). These two spaces are considered naïve because they make no assumptions about underlying developmental processes (Fig. 1Bii). As an evo-devo inspired space (Fig. 1Bi), we used the ICM description of the molar development^35^ and constructed a morphospace based on the relation between the relative occlusal area of m2 and m3 in relation to m1 (m2/m1 and m3/m1, respectively).

We performed model-based clustering analyses of each set of measurements to test our prediction that development will generate discontinuous morphospaces. If, as explained above, complex developmental processes generate a patchy and discontinuous morphospace, then we expect to evaluate a high number of clusters on naïve spaces. However, if the evo-devo informed space corrects this issue, we will observe fewer clusters on the ICM morphospace. As expected, our clustering analysis shows a tendency of the ICM space to find fewer groups than the naïve spaces, suggesting the former is less patchy than the latter (Fig. 2).

**Figure 2.**
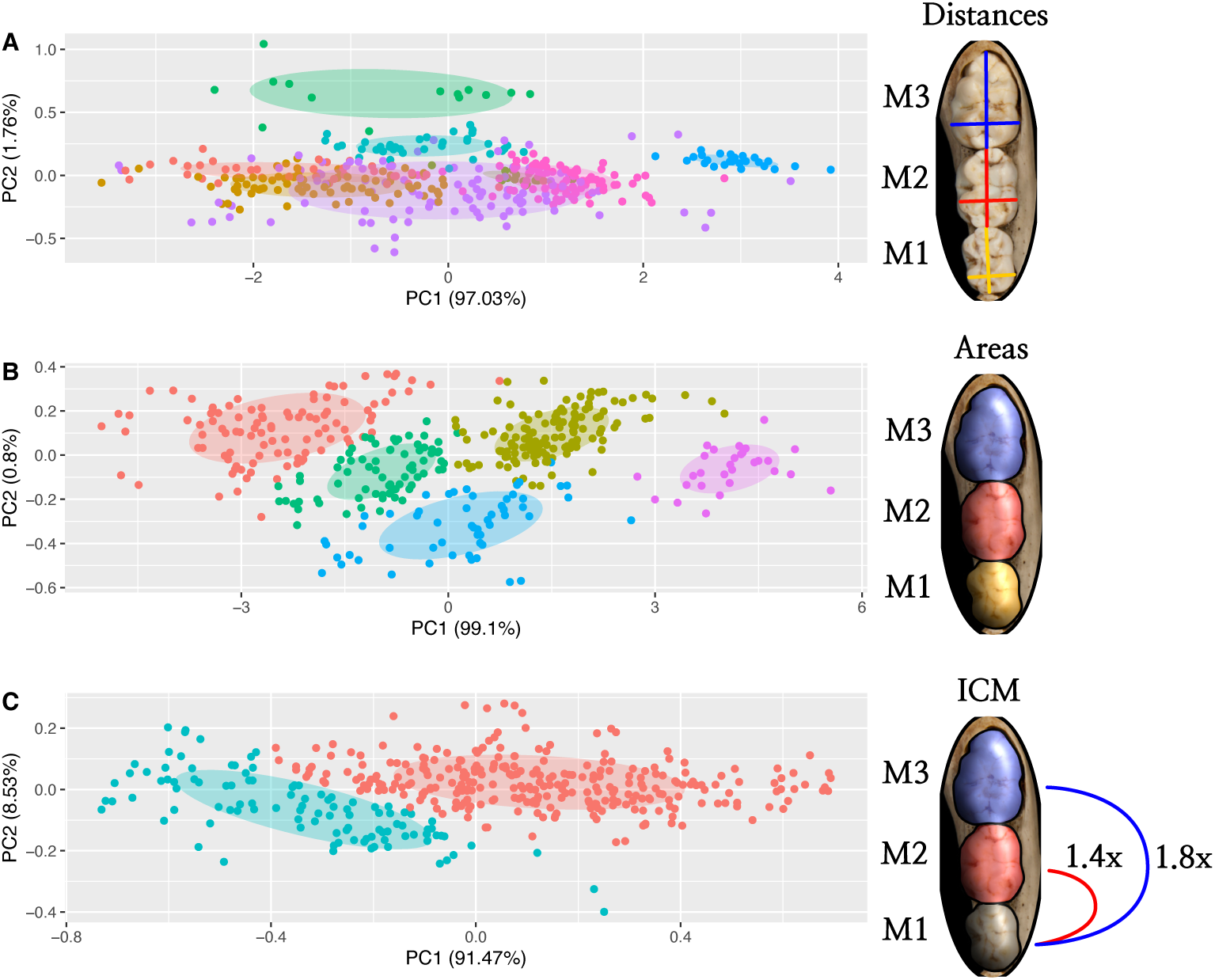
Principal component analysis of the full-sample covariance matrix for the linear distances (A), areas (B) and ICM ratios (C) morphospaces. Dots represent species averages, and colors represents groups identified in the clustering analysis. These groups are based only on morphometric proximity and do not represent any taxonomic group. The more groups a morphospace has, the more patchy and discontinuous it is considered. Axes are not depicted to scale for convenience, so distances in the graph should not be considered representative of the metric of the underlying space.

Our model-fitting approach showed that the use of biologically naïve morphospaces favors evolutionary model complexity. Specifically, the best model for these spaces was a multi-regime multivariate Brownian motion (BM) model (Table 1, S1,S2). BM is a stochastic model in which divergence accumulates linearly with time and is associated with genetic drift under a strictly microevolutionary interpretation or random selection under a macroevolutionary interpretation. For this model, the main parameter is the rate matrix Σ, which controls the traits’ stochastic rate of evolution. Because the preferred model was a multi-regime one, our tree is subdivided into different “regimes,” which are parts of the tree with different model parameters (rates of evolution for BM). For molar occlusal areas, the best model had three main regimes (Fig 3). The first regime covers most fossil groups (thus named “ancestral regime”), including Plesiadapiformes, stem-Haplorhini, part of stem-Simiiformes, and Tarsiidae. The second regime refers to Strepsirrhini, both crown and stem, and the third refers to crown Simiiformes (monkeys and apes, including humans). This three-regime model was also considered a better fit than any global model (microevolution-inspired or not) for the morphospace defined by linear distances, even though it was not the best solution found (see Supplementary Information, Table S2). In both morphospaces, the ancestral regime accumulated more variance over time than any derived regime, suggesting a weaker constraint on the former (Fig S8). It may be tempting to assign interpretations that are either biological (e.g., higher divergence rates after the K-Pg extinction event resulting from ecological opportunity) or statistical in nature (e.g., increased phylogenetic uncertainty of fossil placement resulting in upwardly biased rates^51^). However, we find that such patterns do not appear universally across morphospaces, and is absence from the developmentally-informed one, thus making any interpretation of these partitions premature.

**Figure 3.**
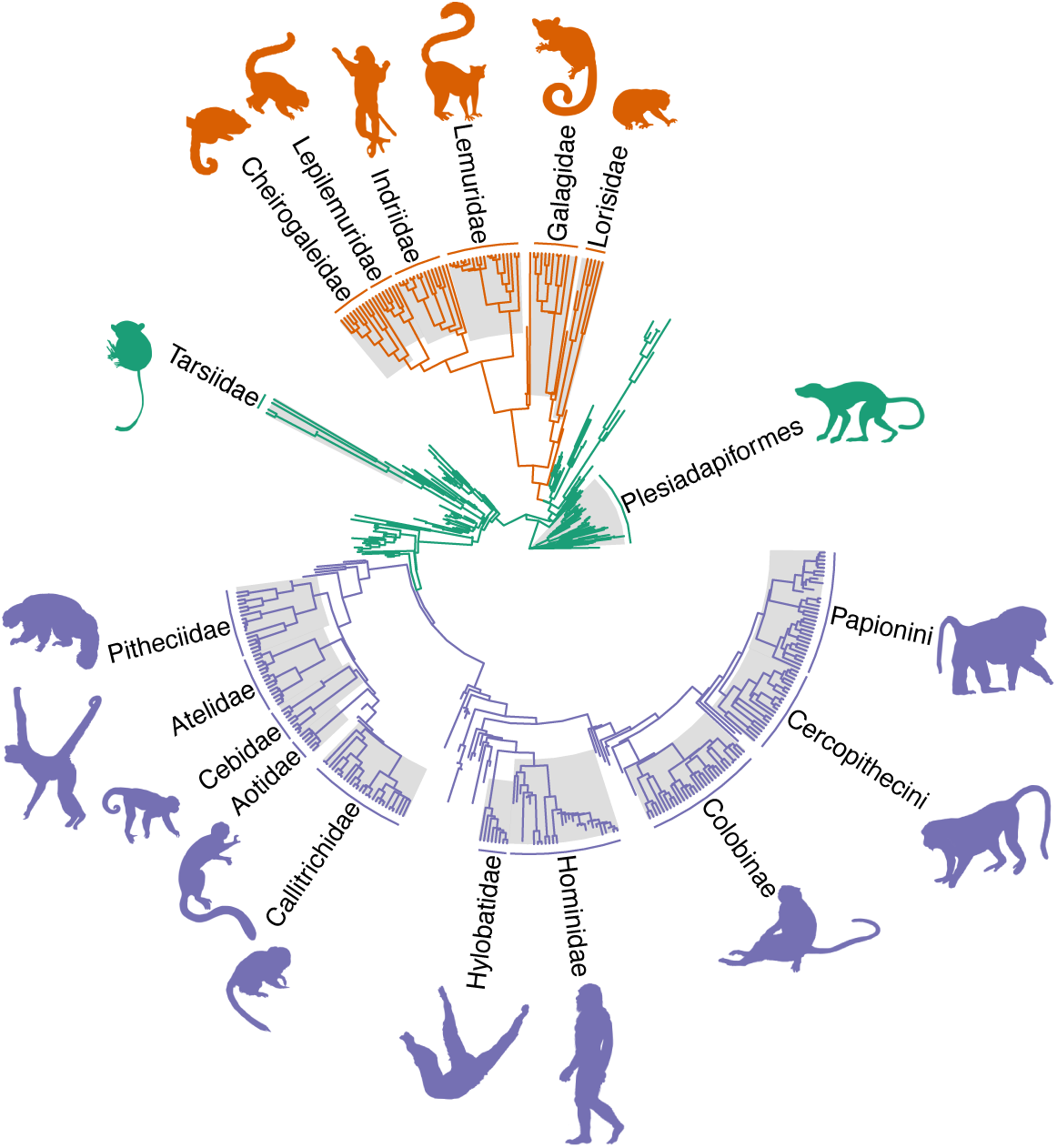
Primate phylogenetic tree painted by the best regime combination found on the phylogenetic mixed model search for the individual molar areas. A multi-regime model allows each different part of the tree (regimes) to have a different model and/or parameter combination. For areas and linear distances, the best model overall is a multi-regime BM, meaning the different highlighted clades will have different rates of stochastic evolution (Σ). For ICM ratios the best mixed model is a multi-regime OU, meaning that each clade will have different rates of stochastic (Σ) and deterministic (*H*) evolution and optima (*θ*). However, for ICM ratios, single-regime microevolutionary models outperform all mixed models (Table 1).

**Table 1.** Model comparison for the primate lower molar row evolution fit through Maximum Likelihood (ML), and ranked according to the BIC.

| Traits<sup>a</sup> | Model<sup>b</sup> | $N_p$<sup>c</sup> | logLik<sup>d</sup> | BIC<sup>e</sup> |
| --- | --- | --- | --- | --- |
| Linear distances | BM | 27 | 2585.03 | -5003.38 |
|  | OU | 54 | 2684.57 | -5035.75 |
| | BM$_{\Sigma \propto P}$ | 7 | 2108.57 | -4173.93 |
| | OU$_{\Sigma \propto P}$ | 34 | 2174.45 | -4138.99 |
| | BM$_{\Sigma \propto G}$ | 7 | 1480.63 | -2918.04 |
| | OU$_{\Sigma \propto G}$ | 34 | 1510.57 | -2811.23 |
|  | Three-regime BM<sup>f</sup> | 69 | 2823.03 | **-5484.50<sup>g</sup>** |
| Areas | BM | 9 | 415.80 | -776.03 |
|  | OU | 18 | 413.98 | -716.83 |
| | BM$_{\Sigma \propto P}$ | 4 | 92.39 | -160.09 |
| | OU$_{\Sigma \propto P}$ | 13 | 275.16 | -470.06 |
| | BM$_{\Sigma \propto G}$ | 4 | 74.41 | -124.13 |
| | OU$_{\Sigma \propto G}$ | 13 | 258.02 | -435.78 |
|  | Three-regime BM<sup>f</sup> | 21 | 471.42 | **-813.19<sup>g</sup>** |
| ICM | BM | 5 | 589.16 | -1147.46 |
|  | OU<sup>D</sup> | 9 | 604.56 | <u>-1153.55<sup>h</sup></u> |
| | BM$_{\Sigma \propto P}$ | 3 | 538.31 | -1058.10 |
| | OU$_{\Sigma \propto P}$ | 8 | 584.68 | -1119.98 |
| | BM$_{\Sigma \propto G}$ | 3 | 575.69 | -1132.86 |
| | OU<sup>D</sup>$_{\Sigma \propto G}$ | 7 | 598.54 | **-1153.86<sup>g</sup>** |
|  | Three-regime OU<sup>f</sup> | 26 | 597.35 | -1034.18 |

<sup>a</sup>Morphospace used to quantify molar form variation; either a biologically naïve space (linear distances or areas) or an evo-devo inspired space (ICM)
<sup>b</sup>Model type. Either global BM or OU models or a mixed model, which allows model and parameter heterogeneity. BM and OU can also incorporate the microevolutionary assumption that the evolutionary rate matrix ( $\Sigma$ ) is proportional to G or P ( $\Sigma \propto G$ or $P$ models). Uppercase "D" indicates OU models with a diagonal $H$
<sup>c</sup>Number of model parameters.
<sup>d</sup>Log-likelihood of the model
<sup>e</sup>Bayesian Information Criterion used for model comparison
<sup>f</sup>Results for the mixed model for Linear distances and ICM are based on the best regime combination found for the areas morphospace
<sup>g</sup>Bold represents the best models
<sup>h</sup>Underline represents the ones with BIC 2 units away from the best model.

By contrast, the investigation of evo-devo-inspired variables based on the ICM paints a strikingly different picture (Fig. 2C). Instead of favoring more complex and heterogeneous models, the ICM morphospace favors a single global Ornstein–Uhlenbeck (OU) process (Table 1). OU models, like BM, have a Σ which governs the stochastic rate of evolution. Differently from BM, however, OU variance does not scale linearly with time, as the evolving species are under the influence of a phenotypic attractor *θ*, to which species converge with a rate governed by the parameter *H*. Under a macroevolutionary interpretation (Fig. 1Di), OU models any evolutionary process with a constraint, be that selective, developmental, genetic, etc. Under a strict microevolutionary interpretation (Fig. 1Dii), Σ is considered the rate of evolution due to random drift, and *θ* and *H* govern the optimum and the shape of the adaptive landscape, respectively. To achieve this interpretation, our microevolutionary model assumes a Σ which is proportional to the additive genetic covariance matrix of the traits (**G**-matrix; equations 1,2, Fig. 1E), with a value within a range governed by empirical estimates of demography and life history of Primates (Fig. 1F-G;^45,46^). This implies that, instead of optimizing values for each entry of Σ, this model only fits one proportionality parameter (*κ*), which makes it simpler and more parsimonious than the macroevolutionary one.

Both macroevolutionary (Fig. 1Bi) and microevolutionary (Fig. 1Bii) versions of this model (OU and OU_Σ∝*G*_, respectively) had essentially the same BICs, suggesting that their information content is effectively the same (Table 1). However, inspecting the confidence intervals for the macroevolutionary OU model reveals that the 95% intervals for its parameters overlap with the values implied by the microevolutionary model (Table S3). This suggests that the OU_Σ∝*G*_ model can be interpreted in terms of microevolutionary processes not only in terms of patterns but also in terms of the magnitude of variation. So, we choose the microevolutionary OU_Σ∝*G*_ as our preferred model not only because it reports the best BIC but also because of its simplicity and biological interpretability.

Following the microevolutionary interpretation of our preferred model, the variation introduced by drift is aligned with the distribution of phenotypes on the ICM morphospace (Fig. 4A), suggesting that the similarity between intra and interspecific patterns of trait variation (see^11^) is consistent with drift. This is further reinforced by the investigation of node-specific rates of evolution, which shows a huge overlap with rates expected under genetic drift (Fig. 5). However, drift alone would generate more variation than the total observed disparity during the period in which Primates have evolved (Fig. S7A), suggesting that stabilizing selection played a crucial role in shaping the pattern of evolution in the group as well.

**Figure 4.**
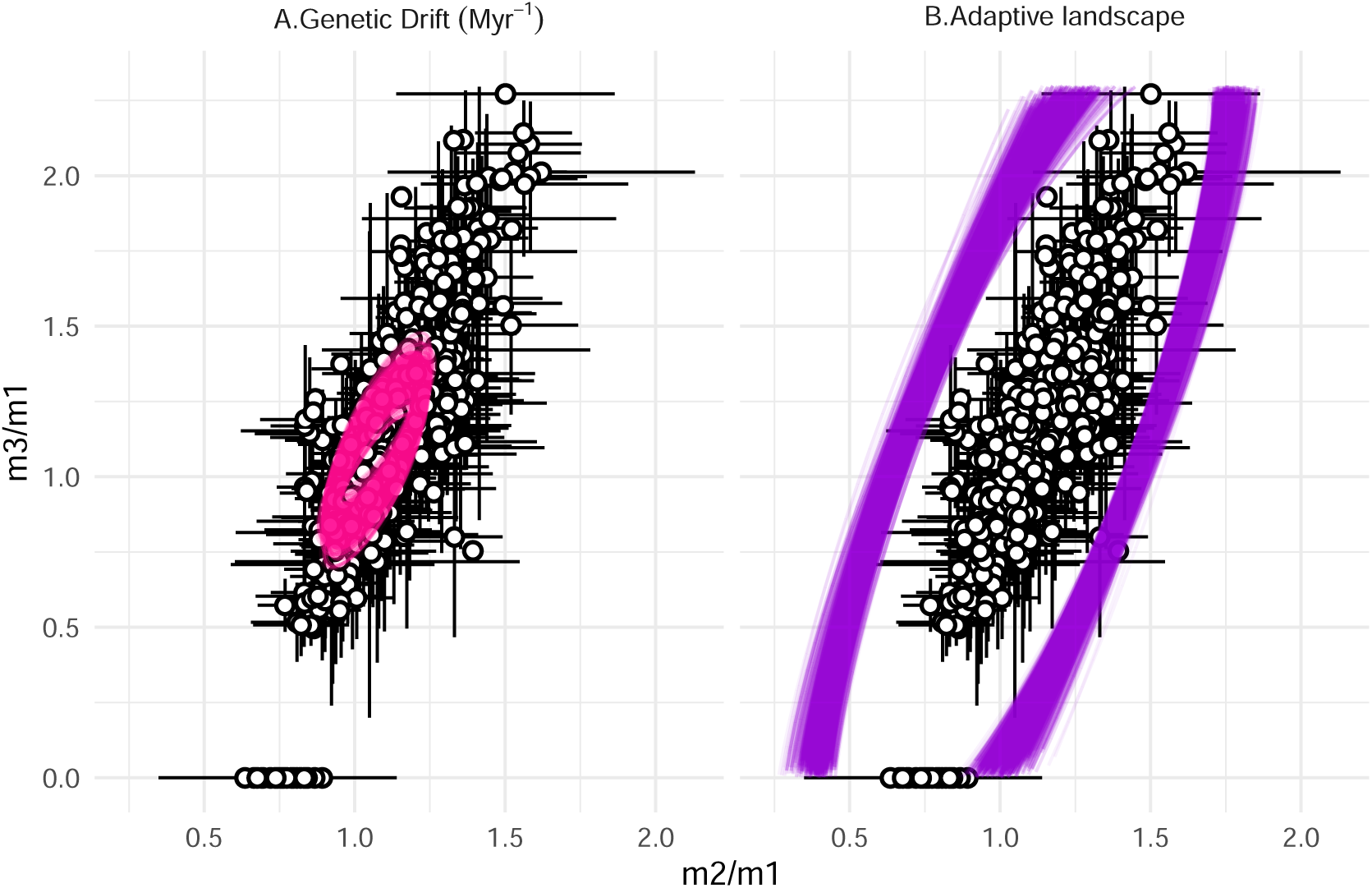
Graphical representation of the best selected model (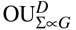) for the ICM variables based on molar ratios m2/m1 and m3/m1. Dots are species averages, and horizontal and vertical lines depict *±*1.96 standard deviations. Ellipses are covariance matrices representing the following parameters of the best model: A. Stochastic rate matrix Σ attributed to the amount of genetic drift introduced in the system every 1 myr. B. Individual adaptive landscape Ω based on model estimates for the rate of adaptation towards the optima (*H*). See Materials and Methods for explanations for these parameters. Multiple ellipses were calculated from parameter value combinations that are sampled along the multi-dimensional likelihood contour 2 log-likelihoods away from the peak.

**Figure 5.**
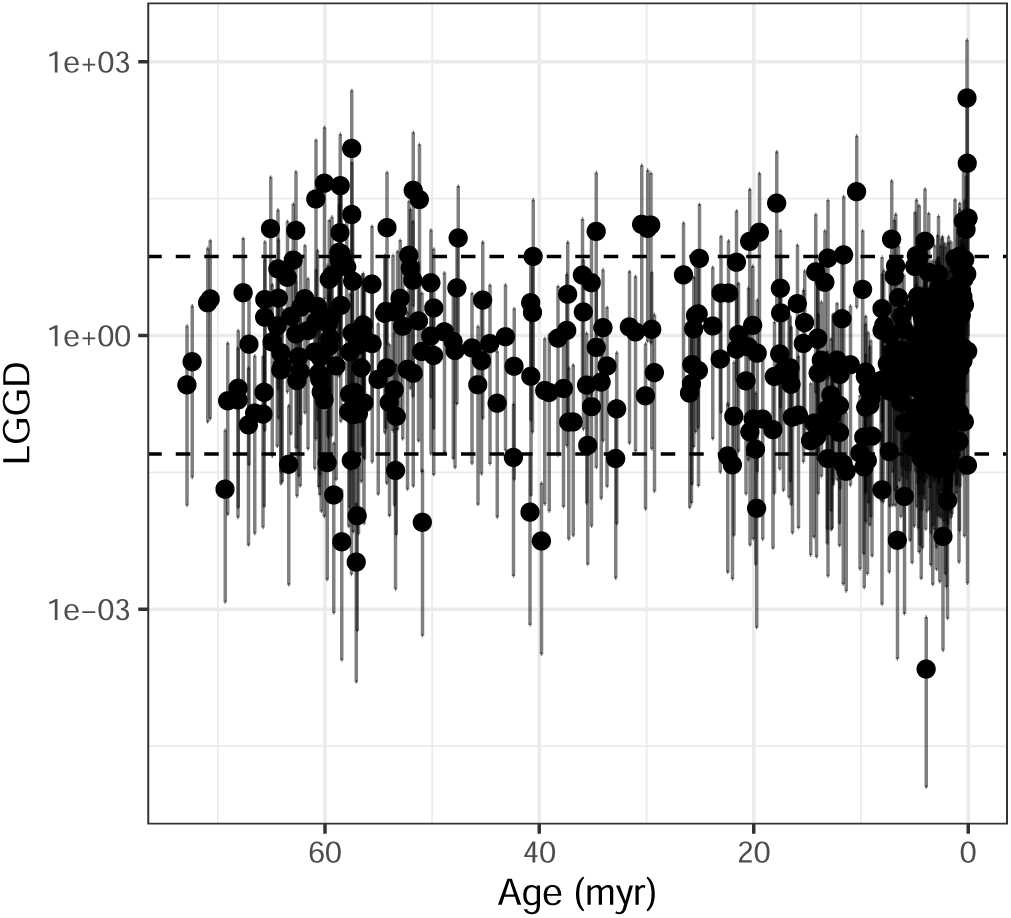
Lande’s Generalized Genetic Distance (LGGD) used to measure the node-specific rate of evolution throughout Primate divergence and diversification. Dashed lines represent the expected rates under genetic drift. For each node, a distribution of values were calculated by integrating over the variation in heritability, effective population size and generation time. Dots represent the median values and vertical lines at the 95% confidence interval.

The investigation of the adaptive landscape implied by the best model shows that stabilizing selection is aligned with the interspecific distribution of phenotypes (Fig. 4B). An examination of the half-lives (*t*_1_*_/_*_2_, the time necessary for a species to reach halfway between the ancestral state and the regime optimum) in different directions of this adaptive landscape shows that *t*_1_*_/_*_2_ are higher along the activation-inhibition gradient direction of the ICM and lower in directions that would lead to deviations from the ICM (Fig. S5). These results indicate that the macroevolution of primate molars is being shaped by a strong stabilizing selection against deviation from the ICM pattern while allowing evolution to occur along the activation-inhibition gradient, in a corridor-like manner.

## Discussion

Previous work has usually highlighted that larger-scale morphological evolution tends to conform to the expectation of microevolutionary models qualitatively but rarely (if ever) in terms of magnitudes of change^6^. In other words, while macroevolution seems to follow directions with more genetic variation, as expected due to neutral change^6,11–13,22,52^, rates of evolution tend to fall below those expected under genetic drift^6–8,53^. This paradox has been used to argue for a fundamental mismatch between micro and macroevolution, as simplistic quantitative genetics models seem unlikely to represent million-year evolutionary processes^7,54,55^. Here, we constrained the proportionality parameter for our preferred microevolutionary-inspired model to be within realistic values for Primates (equation 2; Fig. 1Dii). This constraint results in an estimated rate matrix compatible with drift around a stationary adaptive peak not only in patterns of trait association but also in magnitude. The key modeling choice that led to this conclusion was the quantification of developmentally-informed traits (Fig. 1Bi), which smoothed out transitions between microevolutionary and macroevolutionary data–defining and identifying a neutral subspace aligned with a conserved developmental process. The resulting morphospace of this modeling choice, the ICM space, lacks discontinuities along the diversity of primate molars (Fig. 2), which likely reduces the need for heterogeneous rates along the phylogenetic tree. Furthermore, by focusing on relative shape changes, which are governed by the balance of inhibitory and activation factors, this space limits the influence of other factors, such as static allometry, possibly allowing a closer match between genetic and macroevolutionary variation.

The differences we observe among morphometric representations might be partly due to how different spaces codify size variation. Both naïve spaces contain size information, while the ICM variables do not. By using m1 size as a scaling factor, the ICM variables still include information regarding allometric variation (technically, they are unscaled versions of the Mosimann shape ratios, see^56^). Both area and distance spaces are log-scaled, meaning they fit a power-law allometric model of variation^57^. Therefore, a higher heterogeneity in size variation in the naïve spaces might favor more complex models, while the same is not true for ICM variables. While this suggests size-correction can smooth out much of the heterogeneity in this case, this appears to derive from the fact that using shape-ratios can provide a means to quantify localized ontogenetic effects^58^. Nevertheless, without actual knowledge of developmental systems, it is hard to know beforehand that shape-ratios will necessarily lead to better conformity between micro and macroevolutionary scales. In fact, depending on the system, raw measurements and shape ratios might produce similar results^8^. Thus, studying a well-understood system such as molar development allows us to piece apart the possible role of ontogenetic models in helping us connect micro to macro scales.

Previous work in Primates has suggested that some traits have evolved with rates consistent with those expected under drift^53,59^, including some dental features^60,61^. These works have largely been focused on hominin species, which could bias interpretations regarding the dental evolution of the whole order. Our results partly agree with these results and extend this phenomenon to the group’s origin (Fig. 5). While at face value, this suggests that drift guided over 70 myrs of dental evolution in Primates, our model-fitting approach tells otherwise. Within the microevolutionary-inspired models, OU models outperformed the BM models (Table 1), suggesting a crucial role of stabilizing selection in shaping macroevolutionary patterns. Considering the amount of variation introduced by drift every Myr (Fig 4A), a purely neutral process would result in overdispersion of tip values and higher phylogenetic signals (Fig S7). Instead, the patterns of stabilizing selection seem to be essential in shaping the ICM pattern by both constraining variation that deviates from the ICM pattern and facilitating evolution along the activation-inhibition gradient (Fig S5,S6).

Even though these two results might seem contradictory –rates of evolution consistent with drift and best model including stabilizing selection– we foresee at least two possibilities of how they both might be true: one has to do with the topography of the inferred adaptive landscape, and the other with the estimates of the evolutionary parameters. Regarding the adaptive landscape, the shape of the landscape implied by the preferred model is almost corridor-like (Fig 4B). If this corridor is relatively smooth internally (no great selection differentials within its limits), this would mean that species are free to explore this landscape neutrally, within the bounds of the corridor. Additionally, because the matrix of additive genetic covariances **G** is aligned with the corridor as well (Fig 4A), this means most neutral changes will happen in accordance with the landscape, and will not result in great stabilizing selection. The other possibility is based on the precision of the rate-parameter estimates. Even though the OU model was the preferred one, estimated rates of evolution for BM models are remarkably similar to the ones of the full model S6–S9). Considering that node-specific rates of evolution are calculated under the assumption of a BM model**^?^**^,62^, this could mean that a dense fossil sample in a comprehensive phylogenetic framework might allow for a good estimation of rates of evolution, even under model violation. Irrespective of which is true (or even if both are), the observation that most evolutionary rates are compatible with drift is a pattern rarely seen for macroevolutionary data^6–8,53^.

Together, these results point to the interplay of genetic variation, selection, and development leading to a homogeneous macroevolutionary process within a defined subspace. It has been argued that selection can mold genetic patterns of trait association and variation^29,63,64^, specifically by altering developmental pathways and genetic interactions^65,66^. Conversely, development has also been argued to impose direct selective pressures (*i.e.* internal selection) by reducing the viability of non-conforming phenotypes^29,67^, which could, in turn, trickle down to the organization of genetic variation. While in the present case we can observe this triple alignment between genetics, ontogeny and selection, its origins are harder to decipher. The ICM was originally described in rodents and later verified in many other Mammalian groups^37–40,68^, suggesting it is the ancestral condition for molar development in the group. In this case, ontogeny is viewed as the organizing factor behind both selective patterns and the organization of genetic variation^11^. Furthermore, this explains the near-neutral quality of primate dental evolution, as conformity to the developmental process would be the main selective pressure on relative tooth sizes^69^. However, some Mammalian groups have been shown to deviate from the predictions of the ICM to different degrees, suggesting that the ontogenetic process itself could be malleable^37,38,68,70,71^. Indeed, it has been argued that molar tooth eruption timing in Primates is shaped by biomechanical demands at different ontogenetic stages^72^, revealing a possible mechanism through which external selection could shape development, and indirectly, the morphology of the molar row.

## Conclusions

To what degree microevolution can be extended to macroevolution is a central question in evolutionary biology^4^. While there is little doubt that the fundamental causes at both levels are the same (*e.g.* selection, drift, mutation), efforts to model the connection have generally failed beyond the qualitative alignment of patterns. When it comes to morphological evolution, the consensus has been overwhelmingly to reject any straightforward connection between both levels, specifically because of the fact that empirical evolutionary rates are orders of magnitude inferior to the ones expected by genetic drift^6,7^. The results presented here reject this consensus, as we show that microevolutionary models can fit well to the data, as long as we choose the proper morphometric representation. Even the relatively simple task of characterizing the multivariate dimensions of three molars poses a large number of choices for measurement^10,35,73,91^. Our results suggest that phenotypic quantification based on evo-devo models maximally narrows the gap between both levels of analysis–and allow for the discovery of the underlying subspaces that both qualitatively and quantitatively align macroevolutionary patterns with microevolutionary processes.

While primate molar seems unique in both the presence of a well-constrained ontogenetic model and abundance of data, other systems might also fit the requirements for the methods described here. The existence of evolutionary stable developmental pathways and modules suggests a long history of similarly stable selective pressures^25,65,74,75^. This makes developmental modules good systems to investigate adaptive landscapes in deep-time^32,58,76^. Furthermore, assuming that these pathways are shaped by natural selection to optimize the generation of adaptive variation^63–65,74^, they are a likely place to identify simple connections between micro and macroevolutionary scales^32,58^. So, other evolutionary stable systems are the probable candidates to verify the connection between scales of organization. Good examples are modules built upon serially homologous structures, like limbs, phalanges and vertebrae^77–79^. For more complex structures formed by the interaction of multiple tissues, it might be harder to devise simple models that sufficiently describe the system ontogeny and variation. However, works that focus on the Mammalian skull, and that used individualized bone measurements have had a good track-record of modeling multivariate evolution of these structures under microevolutionary models^12,22,59,80,81^, going even further than the simple alignment between variation and evolutionary rates^8,53,59^. Since vertebrate skull bones are elements with deep individualized history, measuring them individually might represent a good first approximation of the multiple morphogenic fields that interact to form the complete structure. This perspective contrasts with the regular practice of constructing morphospaces as comprehensive, phenomenological, and statistical descriptors of biological form without a clear connection to underlying biological processes^19,36,82,83^. However, given that different morphometric methods seem to point to similar overall patterns of trait variation^84^, finding the correct quantification protocol might be a matter of proper scaling of morphometric variables than a radical departure from classically established measurement practices.

Our investigation also provides a new framework in which developmental biology can be more fully incorporated into macroevolutionary modeling (Fig. 1). It has long been considered that developmental biology was left out of the evolutionary synthesis^26,27,58^, and indeed such data are rarely incorporated into comparative analyses. Recent efforts have had different degrees of success, with many pointing out how complexities of the ontogenetic systems can lead to core violations of the modern synthesis^26,30,31,58,85^. By reframing the question of microevolutionary model adequacy into a problem of quantification of biological phenomena^82,83^, we show how evo-devo is essential for a fully unified view in the context of the evolutionary synthesis.

## Methods

### Sample and morphometrics

We used the standard mesiodistal length (MD) and buccolingual breadth (BL) as basic descriptors of each molar. MD and BL were obtained for each tooth of the lower molar row (m1, m2 and m3). We obtained raw measurements from available datasets in the literature (n=6142)^47–49,97–389^ and from newly measured museum specimens using a caliper (n=150). For rare species, we took measurements from images that were either published or provided to us (n=266). All photos used had a scale and were digitized using the Fiji software^86^. Only adult and not heavily worn teeth where used in our sample, and each specimen was measured once. See the supplementary material for a full list of references and sources used for data collection. In total, we compiled a sample of 6558 individuals distributed among 480 species, divided between 232 extant and 248 extinct species. To evaluate the evolution of these traits on a phylogenetic framework, we used the most comprehensive phylogeny available that included both living and fossil primate species^50^. Our sample covered all genera and 52.98 % of the species diversity included in^50^, spanning the full 75 mya of the group’s evolution.

We constructed three distinct morphospaces to quantify molar variation (Fig. S1). For our biologically naïve representation of tooth form, we used a “distance space” based on linear distances obtained from each tooth and an “area space” based on each tooth’s occlusal area. Occlusal molar area was approximated using a crown index (BLxMD)^38,68^. Both areas and distances where log-transformed to normalize the data and reduce the effect of large-sized outliers. For our evo-devo-informed space, we used the ratios of areas of the second and third molars in relation to the third (m2/m1 and m3/m1, respectively), as defined by the inhibitory cascade model of molar development^35^. We call this last morphospace the “ICM space”. On each morphospace, we calculated species averages for comparative analyses. Measurement error was accounted for by calculating the standard error of each measurement for each species. When a species had a sample of n=1, we assigned a standard error equal to the pooled within-group standard deviation calculated for all species with sample sizes larger than 30. This implies a very high measurement error for species known from single specimens, such as the case of many fossils. The degree of genetic association between traits was approximated both by the intraspecific pooled phenotypic covariance matrix *P*, and an independently derived additive genetic covariance matrix *G* obtained from a pedigreed *Papio hamadryas* baboon population^41,42^.

To evaluate morphospace patchiness we performed a clustering based on parameterized finite Gaussian mixture models (GMM)^87^. This method tests for a series of nested models, where groups are modeled as belonging to different multivariate normal distributions with different group averages. Models differ in the treatment of covariance structures. For example, covariance matrix of different groups might differ in their volume (trace), shape (proportion of eigenvalues), orientation (direction of eigenvectors). Furthermore, covariance matrices might be either spherical (zero covariances, equal variances), diagonal (zero covariances, different variances) or ellipsoidal (non-zero covariances). In total the method tests 14 different covariance models and finds the best partition of the data and the best covariance models according to the Bayesian Information Criterion (BIC).

### Phylogenetic comparative methods

To model morphological evolution, we used a maximum-likelihood (ML) model selection approach, which fits different Brownian-motion (BM) and Ornstein-Uhlenbeck (OU) models under a mixed Gaussian phylogenetic models (MGPM) framework implemented under the R packages PCMbase and PCMfit^88^. This method share some similarities with the GMM clustering method used above to measure morphospace patchiness. Both GMM and MGPM model the data and allow different groups to have different parameter values. However, while GMM fits the data to normal distributions in a non-phylogenetic context, MGPM fits the data according to evolutionary models along a phylogenetic tree. In other words, while the GMM is a non-phylogenetic clustering methods based on species phenotypic proximity in the morphospaces, MGPM groups species according to shared evolutionary models and phylogenetic history.

Under the MGPM framework, the evolution of a p-dimensional multivariate trait is modeled as an OU process as follows:

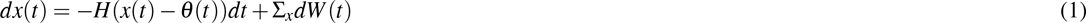

where *H* is the *pxp* selective rate matrix, *x*(*t*) is a *p* vector of trait values at time *t*, *θ* (*t*) is a *p* vector of trait evolutionary optima at time *t*, Σ*_x_* is the Cholesky factor of the *pxp*stochastic rate matrix Σ (sometimes called evolutionary rate matrix) and *W* (*t*) denotes the p-dimensional standard Wiener process.

Under a strict quantitative genetics interpretation^89^, the diagonal of *H* contains the rate of adaptation to the optima of each trait (*α_p_*) and the off-diagonal measures the shape of co-selection among traits. Conversely, the diagonal of Σ contains the rate of evolution due to drift, with its off-diagonal elements containing the amount of coevolution due to genetic covariation. If *H* is a matrix of zeros, the model collapses into a multivariate BM model.

Under this microevolutionary perspective, Σ is not an entirely free parameter. Instead, if Σ is the genetic drift parameter, then it has to be proportional to the additive genetic covariance matrix *G* of those traits^90^ as follows

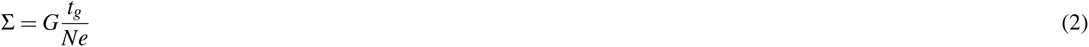

where *t_g_* is the time in generations and *Ne* is the effective population size. Because *t_g_* and *Ne*, and even the size of *G* are hard to estimate at evolutionary time scales, some have argued for treating *t_g_/Ne* as a nuisance parameter, reducing the investigation of drift at the macroevolutionary scale to a simple evaluation of the proportionality between Σ and *G*^12,22,54^. Consistent with these suggestions, here we implement a series of proportionality models, or *κ*-models^11^, in which Σ is set to be equal to a target matrix times a scaling factor *κ*. We used both the intraspecific pooled phenotypic covariance matrix *P* and *G* as target matrices. These models are implemented in the package PCMkappa (https://github.com/FabioLugar/PCMkappa).

Because the proportionality models are more tightly connected to a microevolutionary interpretation of the OU model, we call them “microevolutionary models”. Full models (models where all parameters are estimated freely) are called “macroevolutionary models” because they do not have explicit microevolutionary assumptions.

We fitted two macroevolutionary BM and OU models and two microevolutionary models, using either *P* or *G* as a target matrix, for both BM and OU, totaling six global models (full BM and OU, *BM*_Σ∝*P*_, *BM*_Σ∝*G*_, *OU*_Σ∝*P*_, *OU*_Σ∝*G*_) for each morphospace. For the OU models, we investigated the confidence intervals of the parameters (see Supplementary Material) to evaluate if the model could be further reduced. Specifically, if the confidence interval of the off-diagonal elements of *H* overlapped with 0, another model was fit, setting *H* to be a diagonal matrix^88^.

In addition, we performed a mixed Gaussian phylogenetic model search, which searches for the combination of regimes, models and model parameters that best fit the data^88^. For both the mixed model search and model comparison, we used the BIC, which minimizes parameter inflation due to large samples and is most appropriate for our model-selection question^91^ (i.e. asymptotically identifying the data-generating process as opposed to minimizing trait prediction error). For the mixed gaussian models, we only fit full BM and OU models, and no *κ*-model due to software restrictions. Therefore, the mixed models are also considered macroevolutionary models. All searches where conducted setting the minimun clade size to be five (5) species.

To ensure that the *κ*-models were compatible with microevolutionary processes, we constrained the *κ* parameter to be within the range of expected values under drift, as expressed in equation 2. Because Σ is given in the tree (myr) scale, we found approximations for *t_g_* and of *Ne* for Primates to infer the expected scaling factor *κ*. *t_g_* was estimated as *t_g_* = 1*myr/g_t_*, where *g_t_*is the average generation time in years obtained from^45^. Because we lack good estimates of *g_t_* for fossil species, we used the phylogenetic average *±* the standard deviation (SD) throughout the phylogeny. This was done by trimming the dataset to only the species with *g_t_* data and obtaining the ancestral value and SD at the base through ML^92^. For *Ne* we used 20,000-1,000,000 as the range of possible values consistent with the genomics estimates for multiple primate species and hypothetical common ancestors^46^. While *g_t_* and *Ne* are expected to vary over the tree, we assumed that the effect of this variation would be at least partially canceled out by the fact that these two quantities are generally inversely related to each other.

To evaluate the fitted model mechanistically under quantitative genetics theory, we generalized the equation for the adaptive landscape^89,93^ to the multivariate case as

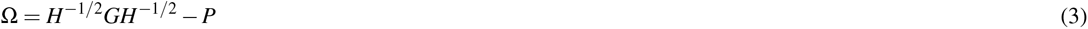

### Rates of evolution

Rates of evolution were used to evaluate if the evolutionary change conforms to the expectation of genetic drift. To calculate rates of evolution, we employed Lande’s generalized genetic distance (LGGD,^90^)

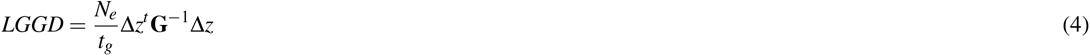

where Δ*z* is the phenotypic divergence calculated as the time-standardized phylogentic independent contrasts (PIC) for each node^8,62^. We produced a distribution of values for each node by sampling values of *G*, *N_e_* and *t_g_* from a uniform distribution between the range defined above. Confidence intervals for the null hypothesis of drift were generated from simulations based on equation 2^8^. Values that fall within the bounds of the null-distribution are thought to conform to the expectation under genetic drift. Values that fall above or below are thought to be indicative of directional or stabilizing selection, respectively.

## A Morphometrics

### A.1 Morphospaces

We used three morphospaces to test our hypothesis that evo-devo-informed quantification would provide a better bridge between micro and macroevolution (Fig S1). The first is based on the linear distances taken directly from the teeth, constituting a 6-dimensional space (“distance space”). The second morphospace is based on the occlusal areas of each molar, which were calculated using a rectangular approximation^38,68^, producing a 3-dimensional space (“area space”). These two spaces are considered naïve because they make no assumptions about underlying developmental processes. As an evo-devo inspire space, we used the ICM description of the molar development^35^, and constructed a morphospace based on the relation between the relative occlusal area of m2 and m3 in relation to m1 (m2/m1 and m3/m1, respectively), resulting in a 2-dimensional space (”ICM space”). Variables were log-transformed in both the trait and area datasets, but not in the ICM space.

To evaluate morphospace patchiness we performed a clustering based on parameterized finite Gaussian mixture models^87^. This method test for a series of nested models, where groups are modeled according to different covariance structures that can be either spherical (all variances equal, no covariances) or ellipsoidal (different variances and non-zero covariances), and can share or not aspects of their covariance matrices, like size, shape or orientation. The permutation of these aspects produces a total of 14 total models, which are fitted with an Expectation-Maximization algorithm, and compared through BIC^87^. In the present case, the best models were the ones in which covariance matrices were ellipsoidal with the same shape, but with either the same orientation and different volumes (VEE) or the same volume and different orientations (EEV). Specifically, both naïve spaces showed a preference to the VEE model, whereas the ICM space showed a preference for the EEV model (Fig. S2). Furthermore, both naïve spaces showed a tendency for finding more groups, with the maximum BIC associated with eight clusters for linear variables and 5 for areas, while the ICM space preferred only two clusters (Fig. S2).

To visualize these results, we performed a Principal Component Analysis (PCA) on the variance-covariance matrix for the full sample on each space. We then inspected the two leading PCs to evaluate the space discontinuity. Because the ICM space is only made up of two variables, the leading two PCs represent all variation on the sample. The inspection of the leading PCs, and the distribution of the groups found through the clustering analysis show that the naïve spaces are more patchy than the ICM space (fig. 2).

To visualize these results, we performed a Principal Component Analysis (PCA) on the variance-covariance matrix for the full sample on each space. We then inspected the two leading PCs to evaluate the space discontinuity. Because the ICM space is only made up of two variables, the leading two PCs represent all variation on the sample. The inspection of the leading PCs, and the distribution of the groups found through the clustering analysis show that the naïve spaces are more patchy than the ICM space (fig. 2).

### A.2 G-matrix

To model the evolution of these traits under quantitative genetics models, we need an estimate of the additive genetic covariance matrix **G** for our molar traits. One common practice in comparative analysis is to use the pooled intraspecific phenotypic covariance matrix **P** as an approximation of **G**^8,12,22,94^. This is justified on the bases of the high similarity between **P** and **G** for morphological traits^11,94^. However, given that **P**s contain also non-genetic information, we also used **G** estimated from a pedigreed population of baboons as a direct model of the Primate **G**^41^.

Because the *G* provided by^41^ was estimated for the raw, untransformed BL and MD measures, we had to transform it in order to match our three morphospaces. We did this through a Monte-Carlo approach, in which we used the published *G* and population means to generate *n* = 300 samples of additive genetic effects. This number of samples was chosen to be compatible with effective sample sizes for the pedigreed population, and not over-represent the accuracy of the estimate^41^. For the linear distance space, we took the generated genetic effects and log-transformed them. For the area space, we log-transformed the product of the effects for each tooth BL and MD. For the ICM space we took the product of the effects for each tooth BL and MD and divided the ones relative to m2 and m3 by m1. We then calculated the maximum-likelihood covariance matrix for the resulting effects for each space. This procedure was done 10,000 times, and the mean covariance matrix was taken as a point estimate of *G* for the lower-dimensionality morphospaces (area and ICM spaces).

To test the validity of this approach we compared the simulation results to the analytic approximation for covariance matrices of products of random variables. Specifically, we investigated if the covariance matrix obtained for the product of the additive effect for each tooth BL and MD (area space) are similar to what would be expected analytically as

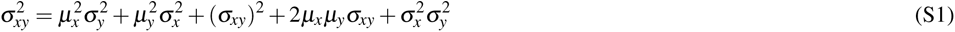

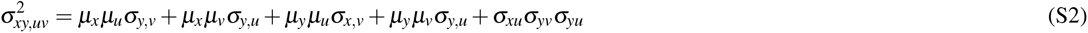

where *µ* are means, *σ* ^2^ variances and *σ* are covariances for the random variables *x*, *y*, *u* and *v*^91^. Variable pairs *x*:*y* and *u*:*v* are BL and MD variable pairs for each tooth. From these equations we were able to construct the covariance matrix for the area space on a *cm*^2^ scale. This procedure was done only for this non-log area space as a proof-of-concept and to limit the number of assumptions necessary to derived log-scale and ratio spaces.

To compare analytical and Monte-Carlo estimates of *G* we first mean-scaled both matrices as follows:

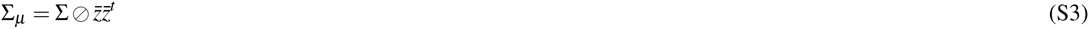

were Σ is the original covariance matrix, *⊘* is the element-wise product and *z̄* is a vector of means^20^. We then calculated the absolute differences from each Monte-Carlo sample of these standardized matrices. Differences in the variances on these matrices are equal to the difference in coefficient of variation between matrices, and thus provide a dimensionless scale of comparison.

The results show that matrices are extremely similar, with a slight bias for covariances being higher on the Monte Carlo samples (fig. S3). Despite this, all absolute differences are very small (*<* 0.002), suggesting that differences are negligible. A matrix correlation analysis reinforced this interpretation, showing values *>* 0.97 for all samples. Together, these results suggest that the Monte Carlo estimates are a reliable approximation for the covariance on lower dimensionalities.

## B Model comparison

We performed model comparison under the maximum likelihood framework of the Phylogenetic mixed Gaussian models^88,92^. This framework has some desirable features. First, it incorporates both measurement errors and missing data during the model fitting procedure. The latter is especially important in our case due to the abundance of fossil taxa in our sample and the presence of species lacking m3s, such as callitrichines and the fossil *Xenothrix mcgregori*. Second, it takes into account process heterogeneity along the tree, allowing regimes to be under different kinds of models and parameter combinations. Lastly, the parameters of the fitted model can be interpreted under quantitative genetics theory in terms of genetic drift and stabilizing selection^55^.

To test our hypothesis, we fit global models that included some microevolutionary assumptions (*κ*-models) and some that did not. Evidence supporting *κ*-models could then be seen as evidence for a microevolutionary interpretation of the data. For each morphometric space, we fitted regular BM and OU models, as well as two *κ* versions of these models. For these models, the rate matrix Σ was set to be proportional to either *P* or *G*. We also performed mixed model heuristic searches, which try to find the best regime combination for a given tree, allowing for regimes to be under different model types (BM or OU). For each space, we conducted 10 searches and recorded the result with the lowest BIC. Lastly, in addition to the heuristic result, we tested mixed models with pre-determinate regime shifts based on Fig. 3 which is the best result for the area morphospace. This regime was chosen as a common point of reference to compare the effect of model and parameter heterogeneity on the fitting process.

### B.1 Distance space

For the distance space only 6 out of 10 heuristic searches converged. Nevertheless, they all produced better BIC values than any global model (table S1). The best model (Search 5) was a multi-rate BM model, with two regimes, one for Simiiformes and one for the remaining of the tree (Fig.S4A). This model deviates from the best model chosen to represent the rate heterogeneity on the other spaces by essentially fusing the ancestral and the Strepsirrhini regimes into one (Fig. 3). In terms of model parameters, this model performed similarly to the tree-regime one in the sense that the ancestral regime had higher rates of evolution than the Simiiformes one (see below fig 5).

Despite the best model being different than the one described in the main text, Search 4 produced a regime combination that was essentially identical to the one for areas (Fig. S4B). Even though the BIC was worse than the best model, this run performed better than any of the global models and is therefore adequate to highlight how this morphospace favors more heterogeneous models instead of global ones.

### B.2 Area space

For the area space, 9 out of the 10 heuristic searches converged. The best mixed model (Search 7) and the best overall model was a multi-rate BM model with three regimes, as described in the main text (Fig. 3).

### B.3 ICM space

For the ICM space, all 10 heuristic searches converged. From these, four runs produced the same result as the global OU model (table S3), suggesting an absence of model shifts for this space. The global fits show that both OU and BM models are equally good fits to the data and that the addition of the microevolutionary assumption of proportionality between Σ and *G* for the global OU model produces a slightly superior model (OU_Σ∝*G*_). Although all three models seem to provide a good fit to the data, the investigation of confidence intervals for parameters of the OU models shows that the confidence interval of the off-diagonal element of *H* overlaps with zero (tables S6,S7), suggesting that this parameter can be excluded from the model (see below).

A model that omits the off-diagonal elements of *H* is implemented as a default model in the PCMfit package^88^. We chose not to include those models initially because our data is rich enough to estimate parameters from even fairly complex models. Instead, we chose to fit the most complex model and perform *post-hoc* model simplification. Using a *κ*-OU model with a diagonal *H* produces a model with vastly superior BIC (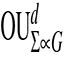). The removal of the off-diagonal elements of *H* did not change significantly the parameter estimates (table S9,S8). Lastly, although OU*^d^* and 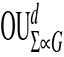 had similar BICs, the confidence interval for all common parameters showed great overlap, suggesting that both models are equivalent. Furthermore, the 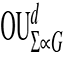 model had tighter confidence intervals, fewer parameters and a slightly larger BIC. For these reasons, we chose this as the best model for our data.

## C Confidence Intervals

Confidence intervals for model parameters and derived statistics were obtained by exploring the likelihood surface and getting parameter combinations from models that were 2 log-likelihood units away from the peak. This was done using the “dentist” package (https://github.com/bomeara/dentist), which allows the exploration of the likelihood surface by “denting” of the surface- setting the region around the peak to have extremely low likelihoods and sampling on the resulting contour around the dented peak. We used 10,000 steps to sample the likelihood surface and obtain confidence intervals simultaneously for all parameters for models fit on the ICM morphospace. If maximum and minimum values sampled by the “denting” approach matched the confidence intervals, then the sampling did not encompass the true confidence interval, and the sampling range had to be expanded. Below we report both the confidence interval as well the extreme values examined to show that adequate confidence intervals are being reported.

For the OU and OU_Σ∝*G*_ models, the off-diagonal elements of the *H*matrix (*H*_1,2_) have confidence intervals that overlap with 0, suggesting that this parameter can be omitted from the model (table S6, S7). The removal of the *H*_1,2_ from the model did not change substantially parameter estimates (table S8, S9), nor did it reduce the likelihoods of the models, improving BIC scores (table S3). All models showed a great overlap in the common parameters (table S6–S9). This is also true for the reconstructed Σ matrix for the *kappa*models. Specifically, for 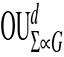, Σ confidence intervals are Σ_1,1_ = 0.0243 *−* 0.0276, Σ_1,2_ = 0.0324 *−* 0.0368 and Σ_2,2_ = 0.0666 *−* 0.0756, placing them within the expected for the OU_Σ∝*G*_ model.

## D Phylogenetic Half-lives

The investigation of the adaptive landscape implied by the best model shows a corridor-like topography (Fig. 4), suggesting that selection is relaxed along the activation-inhibition axis and intensified against deviations from the ICM. To further verify this without assuming a microevolutionary model, we calculated the phylogenetic half-lives (*t*_1_*_/_*_2_) along the activation-inhibition gradient and the deviations from the ICM. *t*_1_*_/_*_2_ measures the time it takes for the phenotype to move halfway in the direction of the optimum and is calculated as

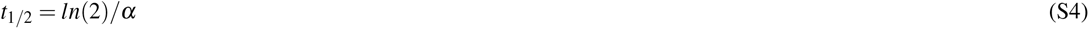

with *α* being the rate of adaptation for each trait given by the diagonal of the *H* matrix^55^. Because *H* is given as a function of the original traits (m2/m1 and m3/m1) we have to perform a rotation of the original matrix into the eigenvectors of the adaptive landscape as

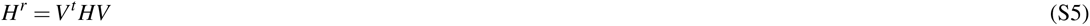

were *V* are the eigenvectors of Ω (see equation 3). *t*_1_*_/_*_2_ were calculated integrating over the confidence limits of parameters. In addition to the rotated space, we also calculated *t*_1_*_/_*_2_ on the original space. This was done to evaluate an additional hypothesis of the ICM process that m3s, because they develop last, are under less intense stabilizing selection^11^.

The results for molar ratios are in line with this prediction, as the half-life for m3/m1 is considerably larger than for m2/m1, suggesting a stronger selection on the latter than in the former (fig S5). For the components of the ICM, results are compatible with the expected for a corridor-like adaptive landscape, with half-lives along the activation-inhibition gradient being higher than on the one for deviations from the ICM (fig S5). This suggests that selection against deviations from ICM is far stronger than the ones along the activation-inhibition gradient, allowing the near-neutral evolution within the adaptive corridor. An inspection of the evolutionary trait-grams of these variables reinforces this idea, as deviations from the ICM are more tightly confined around the optima, and evolution along the activation-inhibition gradient seems greater in amplitude (fig S6).

## E Disparity and phylogenetic signal

To illustrate the effect of stabilizing selection on both overall disparities in the sample and phylogenetic signal, we employed a simulation approach. We generated tip data using the phylogenetic tree and the best evolutionary model in two situations. In one, we used the whole model to produce data under an OU model; in another, we set the *H* matrix to be 0, producing a BM model with the same rate parameters as the OU model. For each model, we generated 1,000 datasets, and for each of these datasets, we extracted the overall disparity and the phylogenetic signal. The disparity was calculated simply as the sum of variances in the tip data, as a measure of overall morphospace occupancy^93^. For Phylogenetic signal, we employed the multivariate version of Blomberg’s K^94,95^. Results from the simulations show the expected pattern, with the BM model showing higher disparity and phylogenetic signal than the OU model (Fig. S7). Because these simulations are based on our preferred microevolutionary model (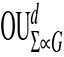), the BM represents what would take place under a pure genetic drift process, while the OU model represents the action of stabilizing selection in modulating the action of drift. So, despite the fact that most of the divergence within the group is compatible with drift (Figure 5), drift alone would produce a wider range of phenotypic values (Fig. S7A, left panel), thus requiring the action of stabilizing selection to not only constrain the total amount of divergence, but also to shape the pattern distribution of phenotypes (Fig. S7A, right panel).

## F Regime-specific disparity

To compare the disparity-generating potential of different regimes of the mixed-model depicted in Fig. 3, we employed a simulation approach. Specifically, we took each model for each regime and simulated phenotypic evolution according to that model on a star-phylogeny of equal size to the full phylogeny. This was done to standardize differences in tree structure, species sample size and model differences (BM or OU) between regimes. Simulations were performed 100 times, and for each run we computed the disparity of the simulated tip data. This was done for the three-regime model for each morphospace. Higher and lower values of disparity indicate a less or more constrained evolution, respectively. Results show that, for all spaces, the ancestral regime is less constrained than the Strepsirrhini and Simiiform regimes (Fig. S8). For areas and distances, the difference between the ancestral and derived regimes disparity is greater than for the ICM morphospace, with Simiiform showing a higher disparity than Strepsirrhini.

## G Supplementary references

## H Data source references

**Figure S1.**
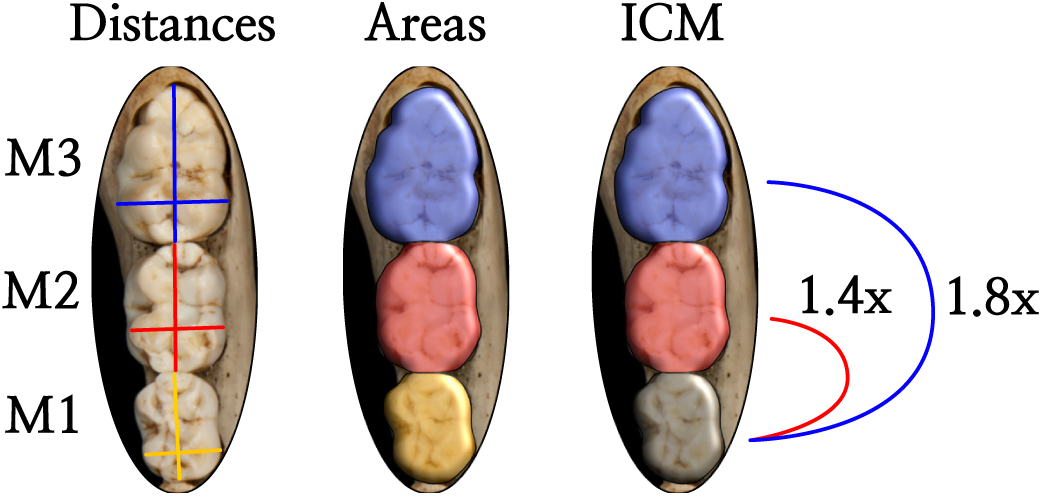
Schematic representation of variables used to construct morphospaces. The trait-space was built on the mesiodistal length (MD, vertical) and buccolingual breadth (BL, horizontal) taken from each molar. The area-space was built by estimating the occlusal areas of each molar as the *A* = *MDxBL*. The ICM-space was built by calculating the relative area size for m2 and m3 in relation to m1.

**Figure S2.**
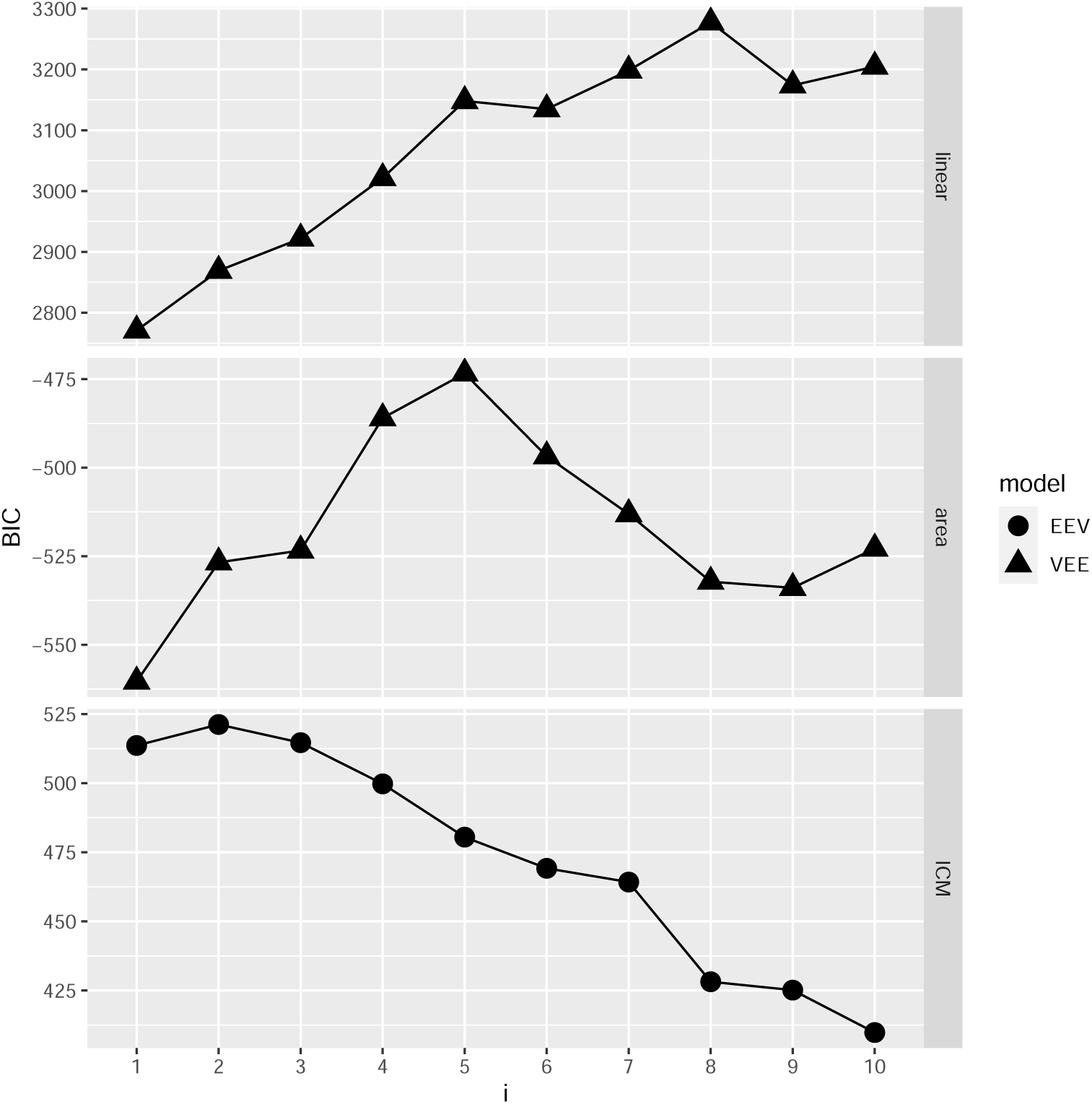
BIC values for the Gaussian mixture models for the linear distances, areas and ICM variables for different numbers of clusters (i). EEV- Elipsoidal model with the same shape, same volume and different orientations VEE- Elipsoidal model with the same shape, different volumes and same orientation.

**Figure S3.**
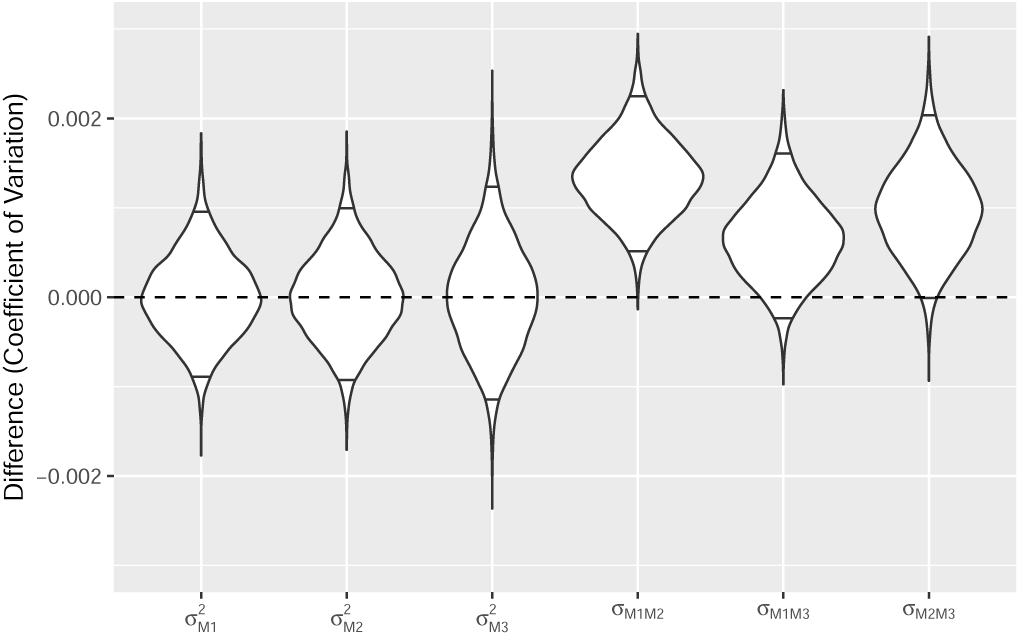
Differences between the Monte Carlo sampling approach for generating covariances for areas and the analytical approximation. Horizontal lines within violins highlights the 95% interval for each matrix cell.

**Figure S4.**
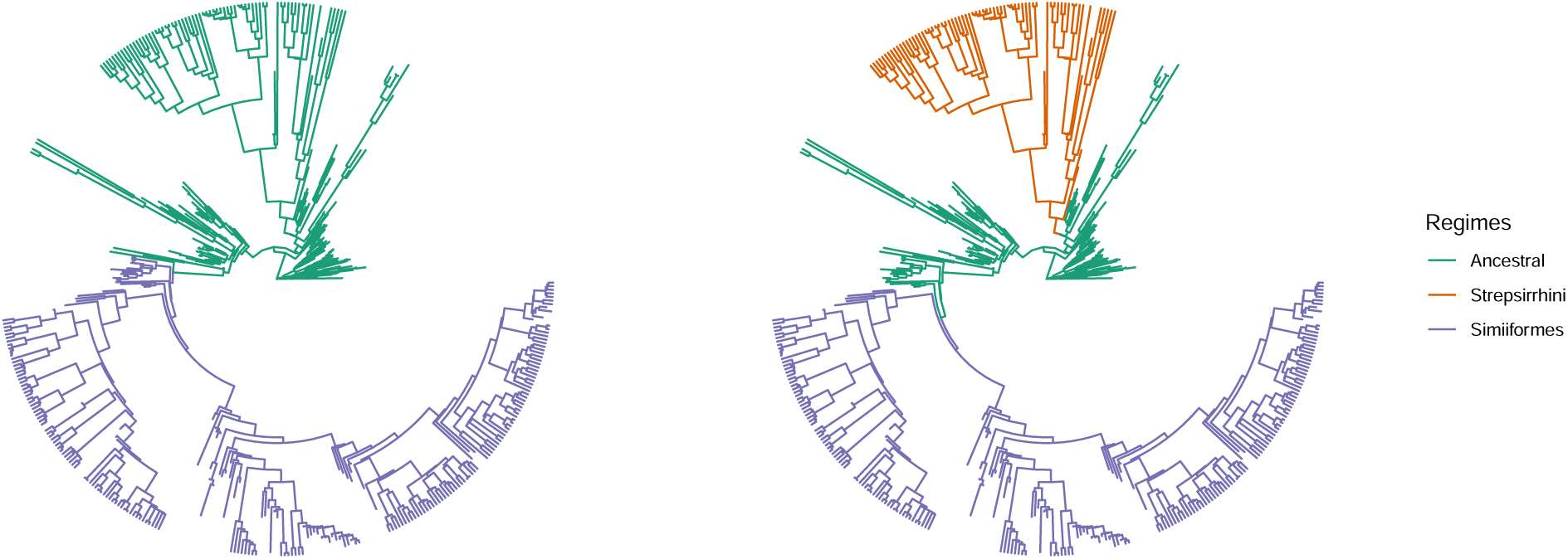
Regimes for different runs of the heuristic search. A- Best model (Search 5). B- Model compatible with the best model for areas (Fig. 3)

**Figure S5.**
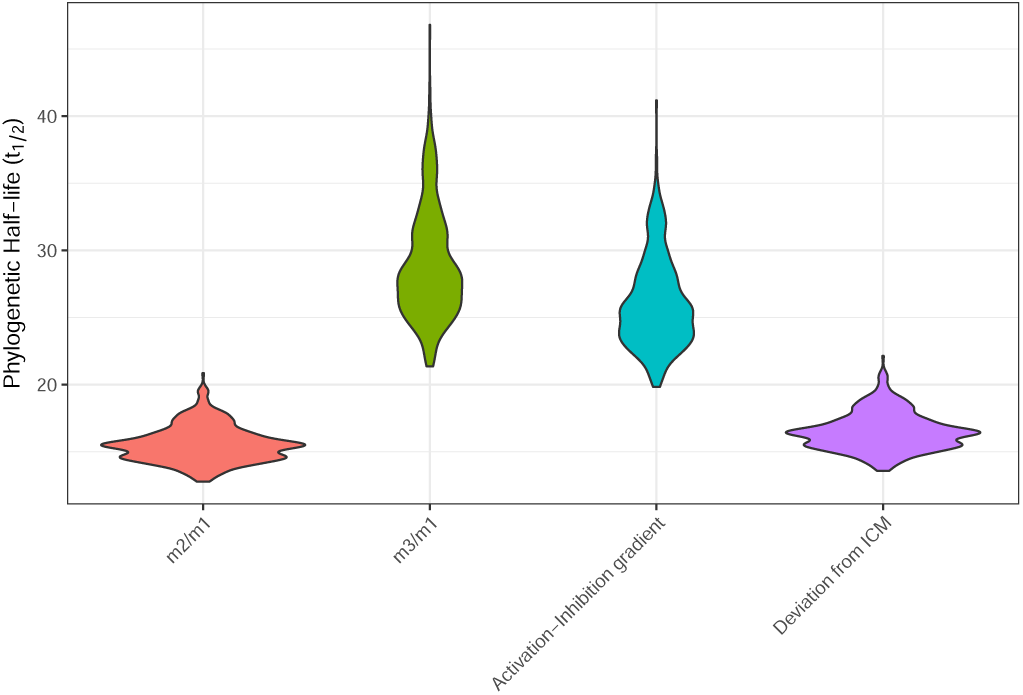
Phylogenetic half life for ICM ratios (m2/m1 and m3/m1) and components of the ICM model (Activation-Inhibition gradient and deviations from the ICM).

**Figure S6.**
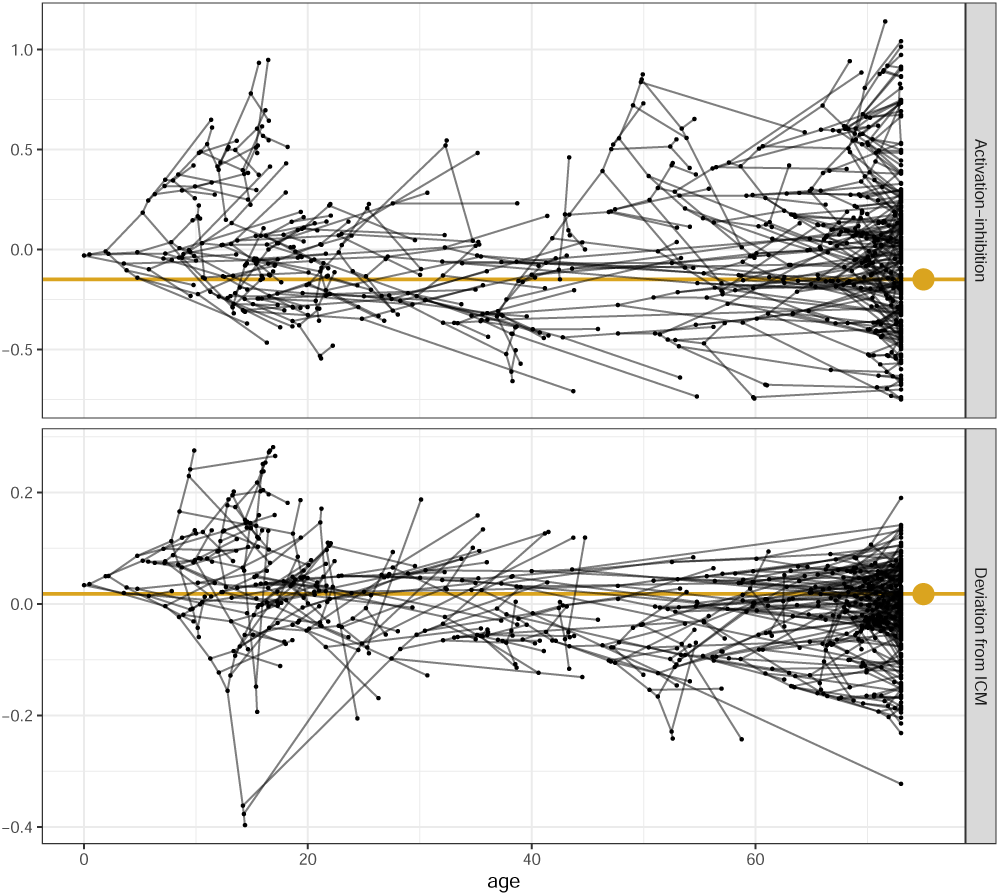
Traitgrams of the components of the ICM for primates. Black lines represents the phylogeny mapped to measured trait values (black points) and golden lines and dots represents each trait optimum.

**Figure S7.**
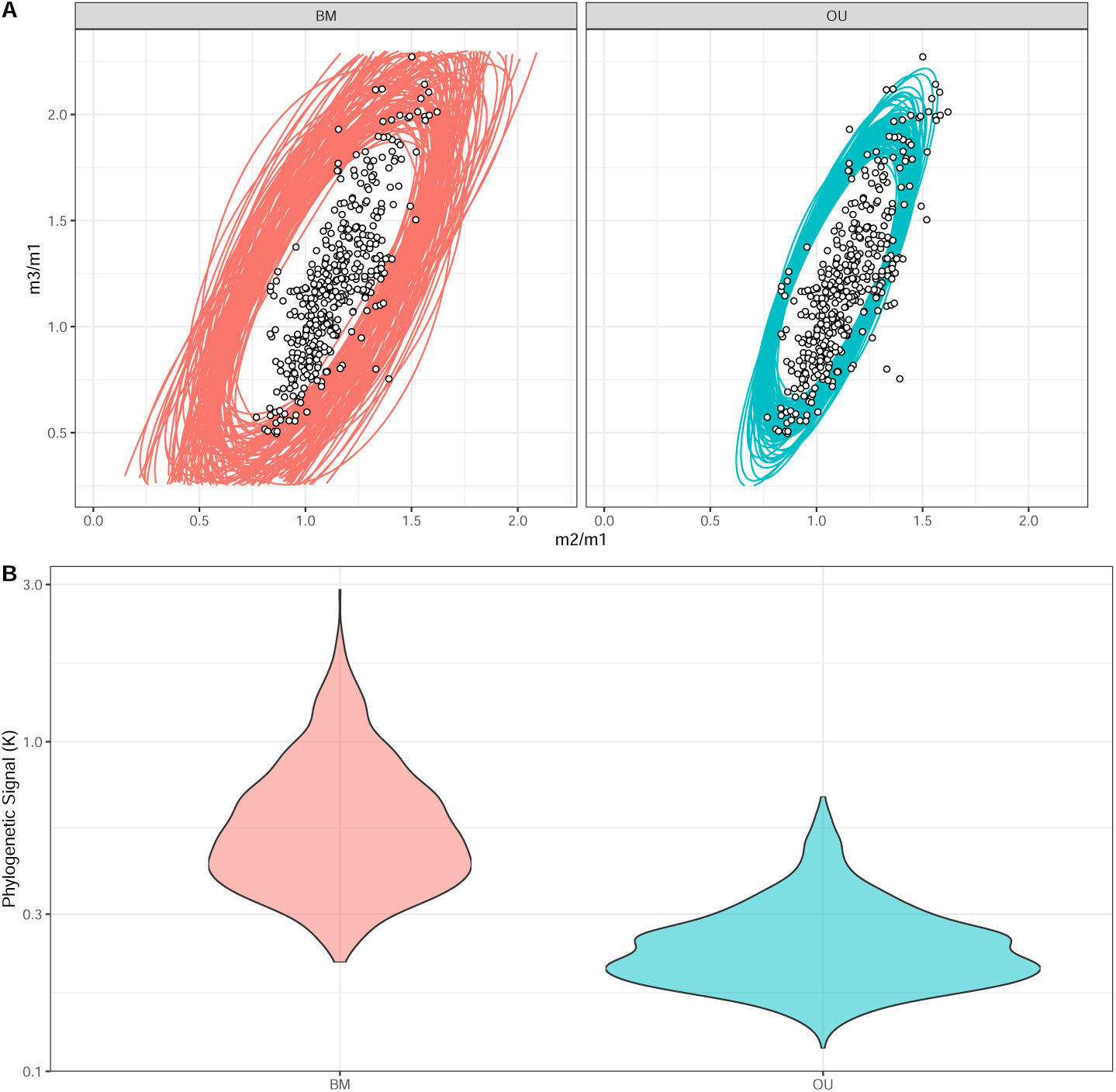
Simulated disparity (A) and phylogenetic signal (B) by assuming the best model (OU) or a brownian motion (BM) model with the same rate parameters as the best OU model. Ellipses represent the covariance matrix of the simulated tip values, and thus do not represent any evolutionary parameter (*e.g.* Σ, *H*, Ω), but the empirical phenotypic distributions. Dots are observed species averages.

**Figure S8.**
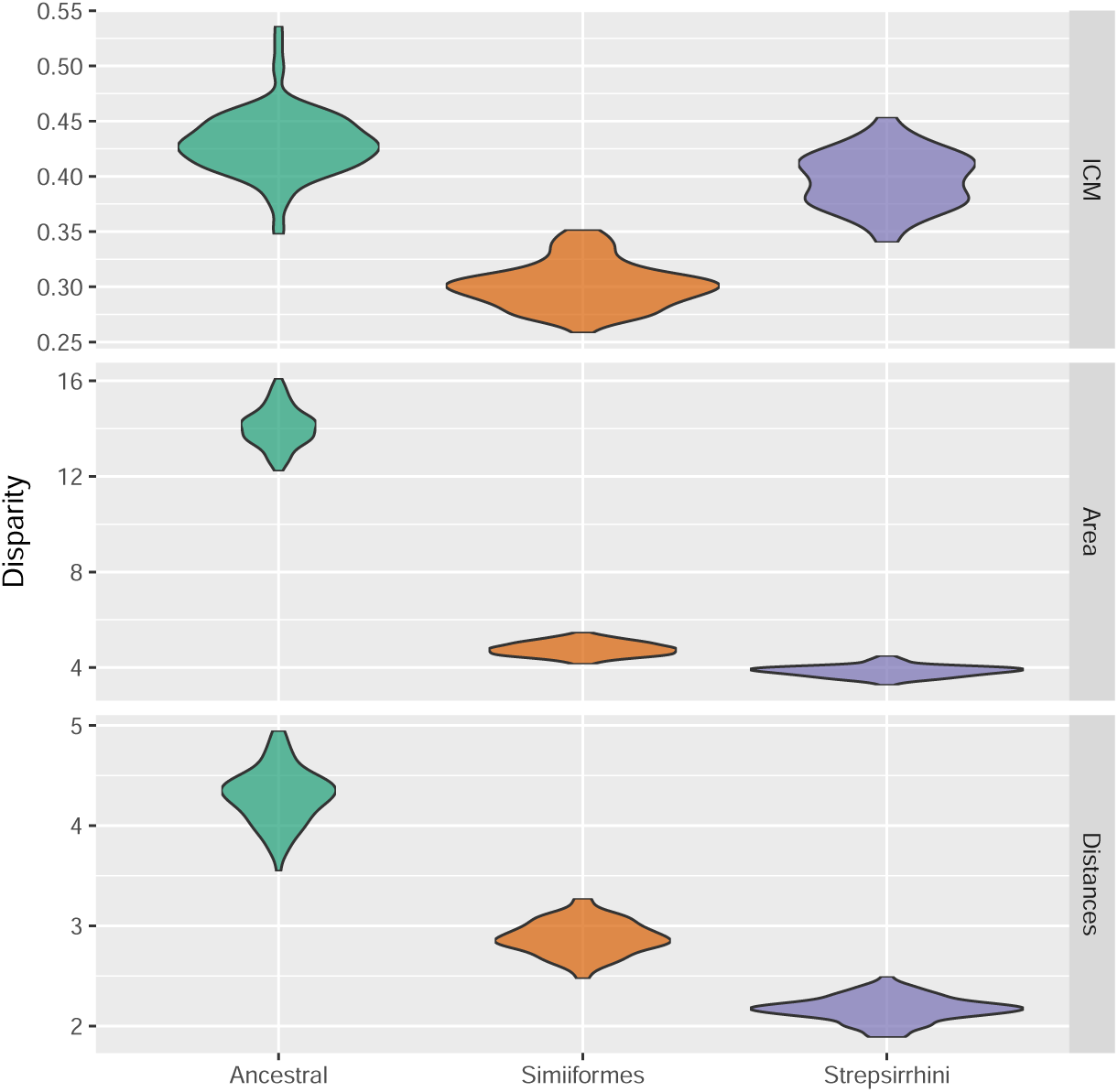
Simulated regime-specific disparities for Three-regime model on morphospace.

**Table S1.**
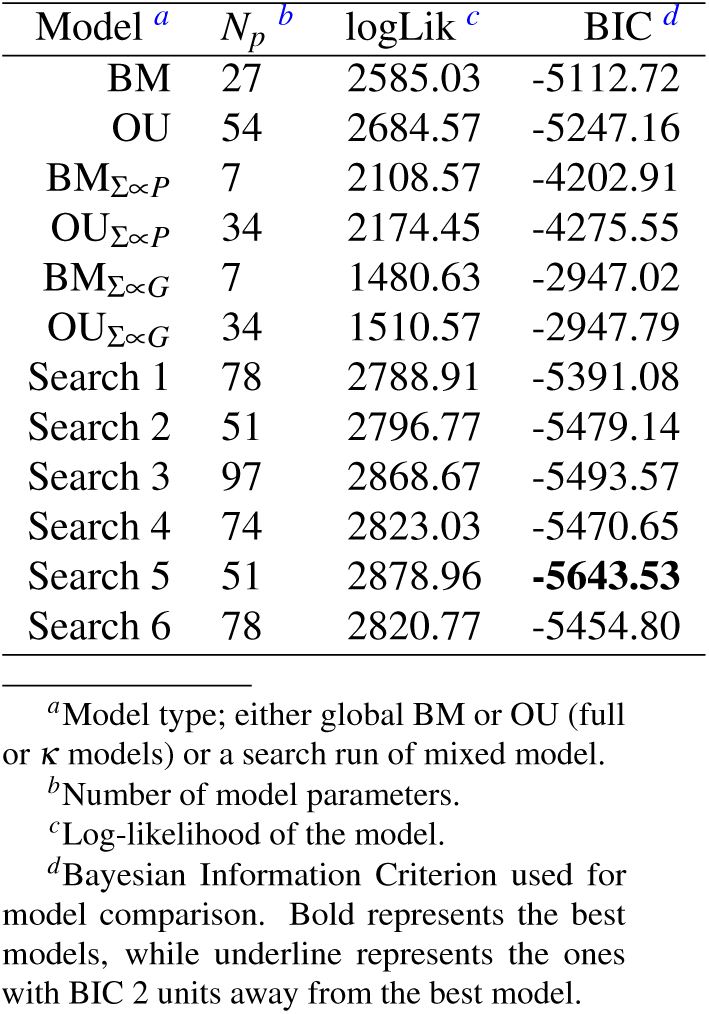
Comparison of models of the linear distance morphospace fit through Maximum Likelihood.

**Table S2.** Comparison of models of the area morphospace fit through Maximum Likelihood.

| Model <sup>a</sup> | $N_p$ <sup>b</sup> | logLik <sup>c</sup> | BIC <sup>d</sup> |
| --- | --- | --- | --- |
| BM | 9 | 415.80 | -813.21 |
| OU | 18 | 413.98 | -790.47 |
| BM<sub>$\Sigma \propto P$</sub> | 4 | 92.39 | -176.70 |
| OU<sub>$\Sigma \propto P$</sub> | 13 | 275.16 | -523.54 |
| BM<sub>$\Sigma \propto G$</sub> | 4 | 74.41 | -140.74 |
| OU<sub>$\Sigma \propto G$</sub> | 13 | 258.02 | -489.26 |
| Search 1 | 10 | 413.61 | -806.75 |
| Search 2 | 10 | 414.98 | -809.48 |
| Search 3 | 19 | 420.25 | -800.85 |
| Search 4 | 10 | 415.85 | -811.23 |
| Search 5 | 18 | 434.43 | -831.37 |
| Search 6 | 26 | 469.34 | -883.58 |
| Search 7 | 26 | 471.42 | **-887.74** |
| Search 8 | 10 | 415.17 | -809.87 |
| Search 9 | 27 | 448.71 | -840.08 |

<sup>a</sup>Model type; either global BM or OU (full or $\kappa$ models) or a search run of mixed model.
<sup>b</sup>Number of model parameters.
<sup>c</sup>Log-likelihood of the model.
<sup>d</sup>Bayesian Information Criterion used for model comparison. Bold represents the best models, while underline represents the ones with BIC 2 units away from the best model.

**Table S3.** Comparison of models of the ICM morphospace fit through Maximum Likelihood.

| Model <sup>a</sup> | $N_p$ <sup>b</sup> | logLik <sup>c</sup> | BIC <sup>d</sup> |
| --- | --- | --- | --- |
| BM | 5 | 589.16 | -1147.46 |
| OU | 10 | 604.53 | -1147.33 |
| OU<sup>d</sup> | 9 | 604.56 | <u>-1153.55</u> |
| BM<sub>$\Sigma \propto P$</sub> | 3 | 538.31 | -1058.10 |
| OU<sub>$\Sigma \propto P$</sub> | 8 | 584.68 | -1119.98 |
| BM<sub>$\Sigma \propto G$</sub> | 3 | 575.69 | -1132.86 |
| OU<sub>$\Sigma \propto G$</sub> | 8 | 598.96 | -1148.53 |
| OU<sub>$\Sigma \propto G$</sub><sup>d</sup> | 7 | 598.54 | **-1153.86** |
| Three-regime OU | 26 | 597.35 | -1034.18 |
| Search 1 | 10 | 604.56 | -1147.37 |
| Search 2 | 15 | 618.60 | -1144.59 |
| Search 3 | 15 | 618.56 | -1144.52 |
| Search 4 | 15 | 617.78 | -1142.95 |
| Search 5 | 10 | 604.55 | -1147.37 |
| Search 6 | 15 | 617.80 | -1142.99 |
| Search 7 | 10 | 604.56 | -1147.37 |
| Search 8 | 6 | 589.16 | -1141.28 |
| Search 9 | 10 | 604.55 | -1147.36 |
| Search 10 | 15 | 617.80 | -1143.00 |

<sup>a</sup>Model type; either global BM or OU (full or $\kappa$ models) or a search run of mixed model.
<sup>b</sup>Number of model parameters.
<sup>c</sup>Log-likelihood of the model.
<sup>d</sup>Bayesian Information Criterion used for model comparison. Bold represents the best models, while underline represents the ones with BIC 2 units away from the best model.

**Table S4.** Confidence interval for the BM model. Lower and upper CI- parameters that are 2 log-likelihood units away from the ML estimate. Lowest and Highest examined values- Extreme values examined by the “denting” approach.

|  | best | lower CI | upper CI | lowest examined | highest examined |
| --- | --- | --- | --- | --- | --- |
| $X0_1^a$ | 1.086 | 0.993 | 1.184 | 0.801 | 1.200 |
| $X0_2^a$ | 1.184 | 1.003 | 1.364 | 0.801 | 1.599 |
| $\Sigma_{1,1}^b$ | 0.018 | 0.016 | 0.020 | 0.010 | 0.499 |
| $\Sigma_{1,2}^b$ | 0.030 | 0.026 | 0.032 | 0.010 | 0.498 |
| $\Sigma_{2,2}^b$ | 0.071 | 0.065 | 0.077 | 0.010 | 0.497 |

<sup>a</sup>Ancestral state at the root
<sup>b</sup>Entries of the rate matrix.

**Table S5.** Confidence interval for the BM_Σ∝*G*_ model. Lower and upper CI- parameters that are 2 log-likelihood units away from the ML estimate. Lowest and Highest examined values- Extreme values examined by the “denting” approach.

|  | best | lower CI | upper CI | lowest examined | highest examined |
| --- | --- | --- | --- | --- | --- |
| $X0_1^a$ | 1.090 | 0.983 | 1.195 | 0.800 | 1.200 |
| $X0_2^b$ | 1.182 | 1.005 | 1.348 | 0.800 | 1.593 |
| $\kappa^b$ | 0.361 | 0.317 | 0.413 | 0.102 | 0.899 |

<sup>a</sup>Ancestral state at the root
<sup>b</sup>Proportionality constant between the target matrix ( $G$ -matrix) and $\Sigma$ .

**Table S6.** Confidence interval for the OU model. Lower and upper CI- parameters that are 2 log-likelihood units away from the ML estimate. Lowest and Highest examined values- Extreme values examined by the “denting” approach.

|  | best | lower CI | upper CI | lowest examined | highest examined |
| --- | --- | --- | --- | --- | --- |
| $X0_1^a$ | 1.132 | 1.008 | 1.200 | 0.801 | 1.200 |
| $X0_2^a$ | 1.306 | 1.075 | 1.466 | 0.804 | 1.597 |
| $H_{1,1}^b$ | 0.034 | 0.025 | 0.043 | 0.010 | 0.498 |
| $H_{1,2}^b$ | -0.214 | -0.485 | 0.473 | -0.490 | 0.498 |
| $H_{2,2}^b$ | 0.031 | 0.021 | 0.040 | 0.010 | 0.498 |
| $\theta_1^c$ | 1.064 | 1.012 | 1.115 | 0.808 | 1.198 |
| $\theta_2^c$ | 1.066 | 0.971 | 1.175 | 0.801 | 1.593 |
| $\Sigma_{1,1}^d$ | 0.022 | 0.020 | 0.025 | 0.010 | 0.495 |
| $\Sigma_{1,2}^d$ | 0.034 | 0.030 | 0.036 | 0.010 | 0.500 |
| $\Sigma_{2,2}^d$ | 0.080 | 0.074 | 0.086 | 0.010 | 0.498 |

<sup>a</sup>Ancestral state at the root
<sup>b</sup>Entries of the **H**-matrix
<sup>c</sup>Multivariate optima
<sup>d</sup>Entries of the rate matrix

**Table S7.** Confidence interval for the OU_Σ∝*G*_ model. Lower and upper CI- parameters that are 2 log-likelihood units away from the ML estimate. Lowest and Highest examined values- Extreme values examined by the “denting” approach.

|  | best | lower CI | upper CI | lowest examined | highest examined |
| --- | --- | --- | --- | --- | --- |
| $X0_1^a$ | 1.097 | 0.966 | 1.192 | 0.800 | 1.199 |
| $X0_2^a$ | 1.232 | 1.031 | 1.376 | 0.802 | 1.599 |
| $\kappa^b$ | 0.486 | 0.416 | 0.553 | 0.101 | 0.894 |
| $H_{1,1}^c$ | 0.049 | 0.037 | 0.057 | 0.010 | 0.499 |
| $H_{1,2}^c$ | 0.226 | -0.352 | 0.469 | -0.497 | 0.500 |
| $H_{2,2}^c$ | 0.024 | 0.013 | 0.035 | 0.010 | 0.499 |
| $\theta_1^d$ | 1.066 | 1.027 | 1.106 | 0.800 | 1.198 |
| $\theta_2^d$ | 1.058 | 0.885 | 1.175 | 0.802 | 1.597 |

<sup>a</sup>Ancestral state at the root
<sup>b</sup>Proportionality constant between the target matrix (**G**-matrix) and $\Sigma$ .
<sup>c</sup>Entries of the **H**-matrix
<sup>d</sup>Multivariate optima

**Table S8.** Confidence interval for the OU*^d^* model. Lower and upper CI- parameters that are 2 log-likelihood units away from the ML estimate. Lowest and Highest examined values- Extreme values examined by the “denting” approach.

|  | best | lower CI | upper CI | lowest examined | highest examined |
| --- | --- | --- | --- | --- | --- |
| $X0_1^a$ | 1.102 | 1.012 | 1.192 | 0.801 | 1.199 |
| $X0_2^a$ | 1.248 | 1.068 | 1.428 | 0.800 | 1.592 |
| $H_{1,1}^b$ | 0.036 | 0.025 | 0.043 | 0.010 | 0.499 |
| $H_{2,2}^b$ | 0.031 | 0.019 | 0.039 | 0.010 | 0.499 |
| $\theta_1^c$ | 1.066 | 1.016 | 1.112 | 0.801 | 1.200 |
| $\theta_2^c$ | 1.073 | 0.969 | 1.211 | 0.808 | 1.585 |
| $\Sigma_{1,1}^d$ | 0.022 | 0.020 | 0.025 | 0.010 | 0.498 |
| $\Sigma_{1,2}^c$ | 0.033 | 0.030 | 0.036 | 0.010 | 0.499 |
| $\Sigma_{2,2}^c$ | 0.079 | 0.073 | 0.084 | 0.010 | 0.499 |

<sup>a</sup>Ancestral state at the root
<sup>b</sup>Entries of the **H**-matrix
<sup>c</sup>Multivariate optima
<sup>d</sup>Entries of the rate matrix

**Table S9.**
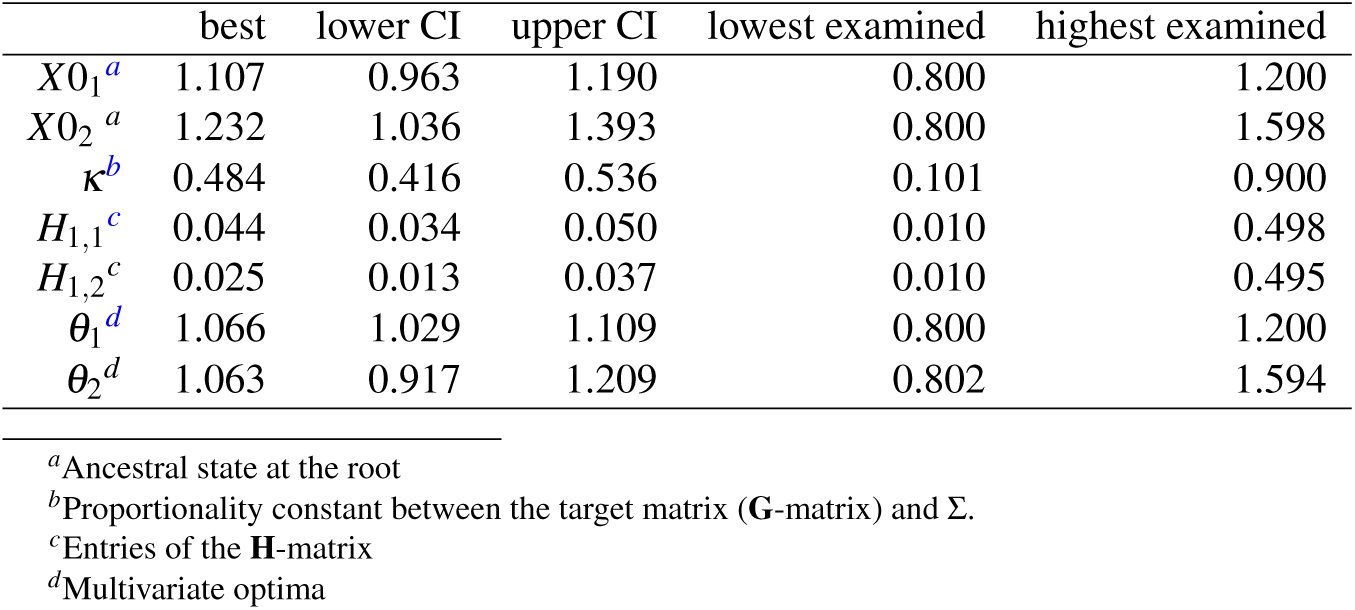
Confidence interval for the 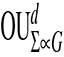 model. Lower and upper CI- parameters that are 2 log-likelihood units away from the ML estimate. Lowest and Highest examined values- Extreme values examined by the “denting” approach.

|  | best | lower CI | upper CI | lowest examined | highest examined |
| --- | --- | --- | --- | --- | --- |
| $X0_1^a$ | 1.107 | 0.963 | 1.190 | 0.800 | 1.200 |
| $X0_2^a$ | 1.232 | 1.036 | 1.393 | 0.800 | 1.598 |
| $\kappa^b$ | 0.484 | 0.416 | 0.536 | 0.101 | 0.900 |
| $H_{1,1}^c$ | 0.044 | 0.034 | 0.050 | 0.010 | 0.498 |
| $H_{1,2}^c$ | 0.025 | 0.013 | 0.037 | 0.010 | 0.495 |
| $\theta_1^d$ | 1.066 | 1.029 | 1.109 | 0.800 | 1.200 |
| $\theta_2^d$ | 1.063 | 0.917 | 1.209 | 0.802 | 1.594 |

<sup>a</sup>Ancestral state at the root
<sup>b</sup>Proportionality constant between the target matrix (**G**-matrix) and $\Sigma$ .
<sup>c</sup>Entries of the **H**-matrix
<sup>d</sup>Multivariate optima

